# Identification of mtROS-sensitive processes in activated CD4^+^ T cells

**DOI:** 10.1101/2020.06.15.152116

**Authors:** Daniel Meston, Wenjie Bi, Tina Rietschel, Marco van Ham, Lars I. Leichert, Lothar Jänsch

## Abstract

T lymphocytes are key components in adaptive immunity and their activation naturally involves mitochondrial-derived oxygen species (mtROS). In particular, H_2_O_2_ has been implicated as an important signaling molecule regulating major T cell functions. H_2_O_2_ targets the oxidation status of functional cysteine residues but knowledge if and where this happens in T cell signaling networks is widely missing. This study aimed to identify mtROS-sensitive processes in activated primary human CD4^+^ T cells. By using a thiol-specific redox proteomic approach we examined the oxidation state of 4784 cysteine-containing peptides of *ex vivo* stimulated T cells from healthy individuals. Upon activation, a shift in oxidation was observed at catalytic cysteine residues of peroxiredoxins (PRDX5 & PRDX6), and T cells were found to maintain their global thiol-redox homeostasis. In parallel, a distinct set of 88 cysteine residues were found to be differentially oxidized upon T cell activation suggesting novel functional thiol switches. In mitochondria, cysteine oxidations selectively modified regulators of respiration (NDUFA2, NDUFA8, and UQCRH) confirming electron leakage from electron transport complexes I and III. The majority of oxidations occurred outside mitochondria and enriched sensitive thiols at regulators of cytoskeleton dynamics (e.g. CYFIP2 and ARPC1B) and known immune functions including the non-receptor tyrosine phosphatase PTPN7. Conversely, cysteine reduction occurred predominantly at transcriptional regulators and sites that coordinate zinc-binding in zinc-finger motifs. Indeed, fluorescence microscopy revealed a colocalization of zinc-rich microenvironments and mitochondria in T cells suggesting mtROS-dependent zinc-release of identified transcriptional regulators including ZFP36, RPL37A and CRIP2. In conclusion, this study complements knowledge on the mtROS signaling network and suggests zinc-dependent thiol switches as a mechanism of how mtROS affects transcription and translation in T cells.

## Introduction

Primary human CD4^+^ T cells are professional mtROS-producing cells capable of utilizing oxidizing species as signaling messengers to elicit physiological immune responses (1). T cell activation by antigen-presenting cells recruits mitochondria to the actin-rich immunological synapse where they support calcium signaling (2) and ATP generation (3). T cells additionally undergo metabolic reprogramming following activation (4) including the increased generation of mitochondrial-derived reactive oxygen species (mtROS) (5). Indeed, T cell activation leads to enhanced electron leakage from complex I (6) and complex III (7) of the electron transport chain (ETC) supercomplex known as the respirasome, resulting in a significant increase in superoxide (8,9). Superoxide is then converted by the mitochondrial superoxide dismutase (SOD2) into hydrogen peroxide H_2_O_2_ (10), which constitutes the major reactive oxygen species in activated T cells and can diffuse to all cellular compartments. On demand, superoxide can also be converted to H_2_O_2_ by zinc-binding superoxide dismutase (SOD1), which is localized in the cytoplasm(11).

Intracellular H_2_O_2_ predominantly targets cysteine residues that are sensitive in their conjugate base thiolate form. Oxidation is mediated by nucleophilic attack of H_2_O_2_ by the cysteine thiolate anion yielding a sulfenic acid which then can conjugate with other cysteine residues to form disulfide bonds (12,13). mtROS-sensitive thiols are those which have a significant fraction existing as the thiolate whereat its stabilization depends on (i) the local ROS and pH level (14) and (ii) the surrounding amino acid residues (15). Therefore, the selectivity of mtROS is dependent on proteins which, by their structure, possess mtROS-sensitive thiolates, ensuring that mtROS only acts on certain redox active protein families. For example, a cysteine which is surrounded with positively charged glutamate residues will stabilize the conjugate thiolate anion, which is then capable of being oxidized (16).

Mechanistically, mtROS signaling can mediate the transient formation and reduction of cysteine disulfide bonds known as thiol switches (17,18). Disulfide bond formation, similar to phosphorylation, affects inter- and intra-molecular interactions and thereby protein functions. A well-accepted example hereof is the regulation of the enzymatic activity of protein tyrosine phosphatases that possess a sensitive thiolate anion in their active site (19) and allow mtROS to regulate phosphorylation signaling (20). Furthermore, quantitative proteomics recently identified redox sensitive thiol switches involved in global translation modulation by mtROS in yeast (21). This involves metal chelating thiolates at zinc-binding proteins (22) and suggested the class of zinc-finger-containing transcriptional regulators as susceptible to mtROS regulation (18). In general, tetrahedrally-coordinated zinc-binding is an evolutionary conserved motif in redox-regulated chaperones (23–25), and zinc-finger proteins in particular have been demonstrated to mediate zinc exclusion and reinsertion through oxidation and reduction of the chelating cysteine (18,21,26,27). With respect to T cells, zinc ions were found to regulate receptor signaling and required to mediate physiological immune responses (28–30). Additionally, T cell activation has been demonstrated to be sensitive to zinc concentration changes by exclusion of free zinc from zinc-binding sites such as in metallothionein (31,32).

The impact of mtROS on the T cell has been studied on both T cell hyper- (33) and hyporesponsiveness (34–36) which suggests diverse and concentration dependent effects on T cell functions. In this line, several reports demonstrated the role of H_2_O_2_ in the fine-tuning T cell metabolism and effector functions (37–40). Additionally, mtROS has long been known to impact T cell apoptosis (41,42) which implicates ROS as a method of immune evasion of tumors (43) and subsequently in health and disease. mtROS is now considered as a mechanistic component of major signaling pathways during T cell activation. This includes the T cell receptor (TCR) signaling pathway, and it was demonstrated that mitochondrial H_2_O_2_ enhances tyrosine phosphorylation of the TCR-associated membrane signaling components Lck, LAT, ZAP70, PLC-γ1, and SLP76 (44). Downstream of the TCR, the nuclear localization of the Nuclear Factor of Activated T cells (NFATc1), which controls T cell colony stimulation and activation, was found regulated by mtROS although the underlying mechanism remains obscure (7,41,45). T cells fine-tune mtROS redox homeostasis for optimal NFATc1 signaling, and higher levels of mtROS in fact downregulate NFATc1 activity in T cells (37). In addition to NFATc1, various other T cell functions such as MAPK signaling (46) as well as PKC-Θ and NF-κB activation (47,48) require an appropriate mtROS homeostasis. As such, a number of antioxidant proteins are present to fine-tune the mtROS signal, including the thioredoxin class of proteins (49), peroxiredoxin family members (50), superoxide dismutase (10,51) and glutathione (52). Peroxiredoxins (PRDX) in particular are known as important in maintaining redox homoeostasis in T cells and are the primary targets of H_2_O_2_ (53). It was demonstrated previously that PRDX regulate T cell expansion and the loss of PRDX2 resulted in immunocompromised mice (54). PRDX also utilize mtROS to elicit a signaling function as has been demonstrated though regulation of the thiol switch tyrosine phosphatase family mentioned previously (55).

In summary, previous studies have characterized how metabolic reprogramming supports mtROS production in activated T cells and demonstrated the importance of antioxidants as part of an regulatory network required for effective immune responses (7,37,48,56). This study now aimed to complement our understanding on where mtROS acts as a signaling component in the T cell proteome. We characterized a superior mtROS inducibility in primary human CD4^+^ T cells as compared to more cytotoxic CD8^+^ T cells and, by using a thiol-specific quantitative proteome approach, identified coinciding redox modifications at cysteine residues. This approach confirmed CD4^+^ T lymphocytes as professional mtROS-producing cells and identified responsive antioxidant peroxiredoxins likely required to maintain global redox homeostasis. In parallel, less than 2 percent of the characterized cysteine residues were found differentially oxidized upon T cell stimulation and revealed a distinct response of the redox proteome. Redox modifications mostly affected functional cysteine residues and were not restricted to the mitochondrial compartment.

## Materials and Methods

### Reagents

The antibodies CD3-BV605 (clone: OKT-3), CD4-PerCP (clone: OKT-4), CD8-PE (clone: RPA-T8) and CD69-PE (clone: FN50) were purchased from Miltenyi (Bergisch Gladbach, Germany). Ficoll was purchased from GE healthcare (Chicago, IL, USA), ultrapure water, DMSO, trifluoroacetic acid (TFA), Tris(2-carboxyethyl)phosphine (TCEP) was purchased form Sigma (MI, USA). All other reagents used were purchased from Thermo Fisher Scientific (MA, USA).

### PBMC isolation

For PBMC isolation and subsequent analyzes of human T cells, Buffy coats from blood donations of healthy human volunteers who provided informed consent were obtained from the Institute for Clinical Transfusion Medicine, Klinikum Braunschweig, Germany. Blood donors’ health is rigorously checked before being admitted for blood donation. This process included a national standardized questionnaire with health questions, an interview with a medical doctor and standardized laboratory tests for (i) infections HIV1/2, HBV, HCV, HEV, Syphilis (serology and/or nucleic acid testing) and (ii) hematological cell counts. 37 ml of peripheral blood obtained from one buffy coat was transferred twice into to 2 50 ml falcon tubes and diluted with 7.5 ml isolation buffer (4% FBS in 1xPBS). Isolated buffy coats were layered over 14 ml of Ficoll separating solution (Sigma Aldrich/Fluka, St. Louis, MO, USA). The tubes were centrifuged at 2000 rpm for 25 minutes (deceleration - 1) at room temperature (RT). Following centrifugation, the PBMC layer was extracted and the cells were spun down at 1600 RPM for 10 minutes (deceleration – 8) at RT and suspended with 50 ml isolation buffer, falcons were spun down and suspended in isolation buffer (as above) twice more. The cell suspension was filtered to remove aggregates and debris (Cell Strainer 40 μm, Falcon). Subsequently, PBMCs were spun down at 1600 RPM for 10 minutes at RT, suspended in RPMI medium (+10% FCS, +1% L-glutamine) at a concentration of 1×10^7^ cells per ml. All subsequent experiments were performed within one day after isolation.

### Fluorescence activated cell sorting (FACS) of CD4^+^ T cells

For cell sorting, PBMCs were spun down at 300xg for 10 minutes at RT and washed in FACS buffer (2% FBS + 2mM EDTA in 1xPBS). Following which, PBMCs were suspended to a cell density of 2×10^8^ per ml in FACS buffer and stained with antibodies against CD3 (BV605), CD8 (PE) and CD4 (PerCP) antibodies at a dilution of 1:50 at 4 ^o^C for 15 minutes. Following which, cells were washed in FACS buffer twice at 300xg for 10 minutes at 4 ^o^C and suspended in FACS buffer. CD3^+^CD4^+^ T cells sorted (Aria-II SORP, BD Biosciences, Franklin Lakes, NJ, USA) with the help of Dr. Lothar Gröbe (HZI, Braunschweig). The gating strategy is depicted in Supplementary Figure 1.

### Immunofluorescence analysis of CD4^+^ T cells for mtROS production

Sorted cells were spun down at 300×g for 10 minutes and cultured overnight in RPMI medium (as above). Cells were first incubated at a density of 1×10^6^ cells/ml in 1 ml RPMI media with mitotracker red (1:10000 dilution) and 5μM H_2_DCFDA in anhydrous DMSO (1:1000 dilution) for 30 minutes in the dark to label the mitochondria as well as load cells with H_2_DCFDA which labels ROS in the cells. Following incubation, cells were washed with PBS before being adhered to poly-L-lysine (Sigma) coated coverslips for 10 minutes. Cells were activated with 10 ng/ml PMA in media added dropwise and images were taken at discrete time periods. Microscopy was performed on an inverted TI eclipse microscope (Nikon,) using a conventional Intensilight epifluorescence Illuminator (Nikon) and data analysis was performed using NIS-Elements AR software by Nikon.

### Immunofluorescence analysis of CD3^+^ T cells for free zinc

Isolated PBMC fractions were resuspended to a concentration of 1×10^6^ cells/ml in 100 μl RPMI complete medium. PMA was added to a final concentration of 10 ng/ml for 1 hour. Twenty minutes before the end of the stimulation, final concentrations of 100 nM MitoTracker Red CMXRos (Invitrogen) and 20 μM Zinpyr-1 (Abcam) were added to the samples. After incubation, 1×10^4^ cells were transferred to a poly-L-lysine (Sigma) coated coverslip and cells were allowed to adhere for another 20 minutes. Then, the supernatant was carefully removed and cells were fixed with 3.7% formaldehyde (Sigma) in PBS for 20 minutes. After washing with PBS and permeabilization for 5 minutes with 0.15% Triton-X100 (Sigma) in PBS, samples were blocked with 1 % BSA (Sigma) and 0.05% Tween20 (Roth) in PBS for 1 hour. Samples were overnight incubated with anti-CD3 (Abcam, clone OKT3, 1:1000) in blocking solution. After three wash steps with 0.05% Tween in PBS, samples were incubated with anti-Mouse Alexa 350 (Molecular Probes, Thermo Scientific, 1:250) in blocking solution for 2 hours. Finally, samples were washed three times, dehydrated with 70% ethanol and 100% ethanol, and mounted using Mowiol (Roth). Microscopy was performed using an inverted Ti eclipse microscope (Nikon), a conventional Intensilight epi-fluorescence Illuminator (Nikon), and data analysis was performed using the NIS-Elements AR software from Nikon.

### Flow cytometric analysis of T cell activation and mtROS production

PBMCs or isolated CD4^+^ T cells were suspended in RPMI media (as above) at a cell density of 1×10^7^ (PBMCs) or 1×10^6^ (CD4^+^ T cells), respectively. Cells were incubated with 5 μM H_2_DCFDA in anhydrous DMSO for 30 minutes and were incubated for 1h with 10 ng/ml PMA at 37 °C. After activation, cells were spun down at 600xg for10 minutes at 4 °C and suspended in 100 μl FACS buffer and incubated with antibodies against CD69 (APC) only in the case of isolated CD4^+^ T cells or, with antibodies against CD69 (APC), CD3 (BV605), CD8 (PE) and CD4 (PerCP) in the case of PBMCs, for 15 minutes at 4 °C in the dark. Cells were washed in FACS buffer before being spun down at 600xg for 10 minutes at 4 °C and suspended in 100 μl FACS buffer for flow cytometry. Analysis of PBMC and isolated CD4^+^ T cell activation and ROS production was carried out on a BD Accuri C6 Flow Cytometer operated with the Accuri CFlow Sampler software (BD, MI, USA). Data analysis was performed using the BD CFlow Analysis Plus software. Additionally, data was acquired on a BD LSR II SORP cytometer (Franklin Lakes, NJ, USA), operated by FACSDiva software. Data analysis was then carried out by using FlowJo software.

### Thiol-selective (iodoTMT) labeling of T cell proteins

To characterize cysteine oxidation in activated T cells, sorted CD4^+^ T cells were spun down at 300xg for 10 minutes at RT and suspended in 1 ml RPMI media (as above) and plated in a 24 well plate at a cell density of 1×10^7^ cells/ml. Cells were either activated with 10 ng/ml PMA or left unstimulated for 1 hour at 37 °C. Cells were washed twice in ice cold PBS and spun down at 300xg for 10 minutes at 4 °C before being transferred to 1.5 ml Lo-Bind Eppendorf cups and lysed in ice cold Denaturing Alkylation Buffer (DAB) (6M urea, .5% SDS w/v, 10mM Ethylenediaminetetraacetic acid (EDTA), 200mM tris-HCl (pH 8.5) +2 μl/ml Benzonase) containing one distinct Thermo Fisher iodoTMT label per condition (active vs unstimulated). The labels used for the first labeling were: 128 for the unstimulated and 129 for the stimulated conditions. Cells were lysed initially for 10 minutes with sonication (high power). Lysates were then incubated at 37 °C with shaking in the dark for 2 hours with shaking in an Eppendorf ThermoMixer allowing all free cysteine residues (R-SH) to be alkylated by the first iodoTMT label. Following initial labeling, the reaction was quenched with the addition of 400 μl of ice-cold acetone and incubated overnight at −20 °C to precipitate protein and remove excess iodoTMT salt. Proteins were pelleted for 30 minutes at 13,000g, −4 °C and washed once with ice cold acetone carefully to not disturb the cell pellet, and excess acetone was allowed to evaporate at RT. Following evaporation, the protein pellet was solubilized with DAB buffer containing 5mM TCEP for 10 minutes at 37 °C to reduce any cysteine residues to a free cysteine (R-SH). Following reduction an iodoTMT label with a different reporter mass for each condition (active vs unstimulated) was added for example 130 for unstimulated and 131 for active conditions, lysates were incubated again at 37 °C for 2h in the dark with shaking on an Eppendorf ThermoMixer. Again, the reaction was quenched with the addition of 400 μl of ice-cold acetone and incubated for 4 hours at −20 °C, to remove excess iodoTMT salt. Proteins were pelleted for 30 minutes at 13,000g −4 °C and washed once with ice cold acetone, and excess acetone was allowed to evaporate at RT. Protein pellet was dissolved in 100 μl Dissolution buffer (0.5M TEAB pH 8.0) and samples were mixed in a 1:1:1:1 ratio. Trypsin was added to a ratio of 1:20 of trypsin: protein, samples were incubated overnight at 37 °C with shaking on an Eppendorf ThermoMixer.

### ZipTip clean-up of peptide samples

Following protein digestion a 5% volume of the peptide solution was cleaned up with a reverse phase (C18) ZipTip (Merck Millipore, Burlington, MA, USA). The peptide sample was acidified with the addition of 1 μl 1M hydrochloric acid. ZipTip pipette tips were equilibrated with 10 μl of SPE cartridge wetting/elution buffer (60% ACN in 0.2% TFA) with 10 uptake and dispensing strokes of the pipette before being equilibrated with 10 μl SPE cartridge washing buffer 3% ACN in 0.2% TFA) with 10 uptake and dispensing strokes of the pipette. Peptide samples were loaded onto the ZipTip with 20 strokes of the pipette. Peptides were washed with 10 μl SPE cartridge washing buffer with 10 slow uptake and dispensing strokes of the pipette, peptides were eluted into a new Eppendorf tube with 10 μl SPE cartridge wetting/elution buffer with 10 fast uptake and dispensing strokes of the pipette. Peptide samples were dried in a SpeedVac (Sorvall/Thermo Fisher Scientific, Waltham, MA, USA) and suspended in 10 μl Nano LC-MS buffer A (0.1% FA) before being injected onto an UltiMate 3000 RSLCnano (Thermo Fisher Scientific/Dionex) coupled to an Orbitrap Velos Pro MS (Thermo Fisher Scientific) to determine digest efficiency as well as peptide amount.

### Solid phase extraction (SPE) of peptide samples

Peptide samples were dried and suspended in 200 μl SPE cartridge washing buffer. Hydrophilic lipophilic balance (HLB) 1CC (10 mg) SPE columns (Waters corporation, Milford, MA, USA) were first wet with 200 μl SPE cartridge wetting/elution buffer (as above) followed by equilibration with 200 μl SPE cartridge washing buffer (as above). Suspended peptides were then loaded onto the cartridge and allowed to flow through the cartridge. Samples were washed with 200 μl SPE cartridge washing buffer, peptides are then eluted with 100 μl SPE cartridge wetting/elution buffer into a 1.5 ml Eppendorf tube, and samples were dried using a SpeedVac.

### Basic reverse phase (bRP) fractionation of peptide samples

Peptide samples were suspended in 300 μl of bRP buffer A (1% ACN in 10 mM Ammonium hydroxide) and loaded into a 500 μl injection loop. Samples were separated on an ÄKTA purifier UPC10 (GE Healthcare, Chicago, IL, USA) consisting of a INV907 column switcher, a M925 UV flow cell coupled to a Frac-950 fraction collector under high pH reverse phase conditions. Fractionation was performed on a Zorbax 300 extend 25 cm x 9.4 mm I.D, 5 μm, 300 Å semi preparative column (Agilent Technologies, CA). The Zorbax 300 column was maintained at 30 °C and a flow rate of 1 ml/min was used. Peptides were separated using a binary solvent system consisting of bRP buffer A and bRP buffer B (90% ACN in 10 mM Ammonium hydroxide). They were eluted with a gradient of 5-28% B in 50 minutes, 26-70% B in 4 maintained at 70% B for 5 minutes and finally the column was re-equilibrated to 5% B in 4 minutes. 1 ml fractions were collected into 2 ml Eppendorf cups every minute. Fractions were dried in a SpeedVac until a small volume remained, following which samples were concatenated to 13 samples by mixing every 12^th^ fraction together. Samples were dried and suspended in 10 μl Nano LC-MS buffer A before being injected onto an UltiMate 3000 RSLCnano coupled to an Orbitrap Fusion MS (Thermo Fisher Scientific) for analysis.

### Nano-LC/MS analysis of iodoTMT peptides

Peptides were separated on an UltiMate 3000 RSLCnano system consisting of a NCF3500PS pump module, a WPS3000 Autosampler and a TCC-3000S column compartment under reversed phase conditions, interfaced via Electrospray ionization (ESI) to an Orbitrap Fusion Tribrid MS. Peptides were washed under non-eluting conditions on a Thermo Fisher Acclaim PepMap 2 cm x 100 μm I.D. 5 μm, 100 Å guard column (Waltham, MA, USA) for 6 minutes, column was maintained at 60 °C with a 1 ml/min flow rate of NanoLC buffer A (0.1% formic acid) before eluting on to the analytical flow path by increase to 6 ml/min flow rate.

Analytical separation was performed using an Acclaim PepMap RSLC 50 cm x 75μm I.D. 2μm, 100 Å column (Thermo Scientific, MA). Column was maintained at 60 °C and a flow rate of 200 nl/min was used. Peptides were separated using a binary solvent system consisting of Nano LC-MS buffer A and Nano LC-MS buffer B (80% ACN in 0.1% FA). They were eluted with a gradient of 3.7–31.3% B in 120 min, 31.3–62.5% in 45 min, 62.5–90% B in 8 min, maintained at 90%B for 5 min and finally re-equilibrated to 3.7% B in 18 min.

Eluting peptides were converted to gaseous ions via a fused silica Picotip 5 cm x 10 μm ID emitter (NewObjective) for ESI (spray voltage, 2 kV; heated capillary, 300 °C) using a nano-electrospray source (Thermo Scientific). MS spectra were acquired in the positive ion mode on the Orbitrap Fusion MS with an m/z range of 375–1,700 with R of 120,000 at m/z 200. AGC target was 4 × 10^5^ with an injection time of 50 ms. The top 50 most intense precursors were selected for higher-energy collisional dissociation (HCD) again on the Orbitrap for high resolution MS^2^ spectra of iodoTMT reporter tags with a collisional energy of 32%. MS^2^ parameters were as follows: charges= 2-4^+^, R= 15,000 at m/z 200; AGC, 5 × 10^4^ with an injection time of 60 ms.

### Identification of iodoTMT labeled peptides

For peptide identification Mascot (57) integrated into proteome discoverer (v2.2 Thermo Fisher Scientific, MA, USA) was used to search peak lists against the H. sapiens proteome (UniprotKB canonical set 2020_02). IodoTMT sixplex reporter tags (126.127725, 127.124760, 128.134433, 129.1311468, 130.141141, 131.138176) were chosen as quantification method. IodoTMT modification (329.226595) was set with cysteine residue specificity, and up to 5 modified amino acids per peptide were allowed. Trypsin/P was set as the enzyme with a specific digest method with up to 1 missed cleavage site allowed. Mass tolerance was set at 7ppm for precursor identification and 50mmu for fragment masses, variable modifications were set as oxidation of methionine and acetylation of N terminal peptides. PSM selection criteria was set at a FDR of 0.01 for only high confidence matches with a minimum peptide length of 6 amino acids, the decoy database was deployed in concatenated mode. The mass spectrometry proteomics data have been deposited to the ProteomeXchange Consortium via the PRIDE (58) partner repository with the dataset identifier PXD019649.

### Protein association network

STRING (59) (version 11.0) was used to generate a functional protein association network using the search form for multiple proteins. The minimum required interaction score was set to highest confidence (0.9). The meaning of network edges was set to the term “confidence” (line thickness indicates strength of data support).

### GO analysis

To analyze the cellular localization of proteins, all quantified cysteine-containing proteins were sorted according to the GO Ontology database terms. Proteins with peptides in multiple oxidation ranges were grouped with the most oxidized peptide. The Panther search function (version 14.0) was used to examine enrichment of GO terms (60). The built-in Bonferroni correction procedure was used to correct raw P-values calculated for GO enrichment analysis for multiple testing. GO terms with a corrected P-value < 0.05 were considered enriched.

### Statistical analysis

Sample size, replicates and statistical tests were performed in accordance with literature with similar methodology. For statistical analysis a two-sided, paired T-test was performed, a p value of 0.05 was taken as the statistical significance minimum, where; * is <0.05, ** is <0.01 and *** is <0.001. All statistical analyses were performed using R studio (61); additionally R studio was used for all data visualization.

## Results

### Mitochondrial ROS induction in CD4^+^ human T cells

In this study, we aimed at identifying cysteine residues that alter their redox status in activated, mtROS-producing T cells. Firstly, we validated the suitability of human primary CD3^+^ T cells as a model to study mtROS generation and concurrent redox proteome. To this effect we utilized the diacylglycerol (DAG) mimetic phorbol myristate acetate (PMA) to induce T cell activation. PMA has been applied previously to study ROS generation (62–64) and it was demonstrated that PMA alone triggers metabolic reprogramming as well as mtROS production in primary T cells (48,65). We applied PMA to total PBMC fractions from healthy individuals to examine the inducibility of mtROS in both major CD3^+^CD4^+^ and CD3^+^CD8^+^ T cell subsets which we comparatively examined by flow cytometry (Supplementary Figure 2). CD3^+^CD4^+^ T cells were found notably more responsive and exhibited a higher dynamic range of ROS induction as compared to the CD3^+^CD8^+^ T cell subset. As such, CD3^+^CD4^+^ T cells were isolated from PBMC fractions of three healthy individuals by using a FACS-based strategy. CD3^+^CD4^+^ T cells were then stimulated *ex vivo* with 10 ng/ml PMA for 1 hour and the activation of T cells could be validated by the surface expression of the early activation marker CD69 in all donors (Figure 1A).

**Figure 1:**
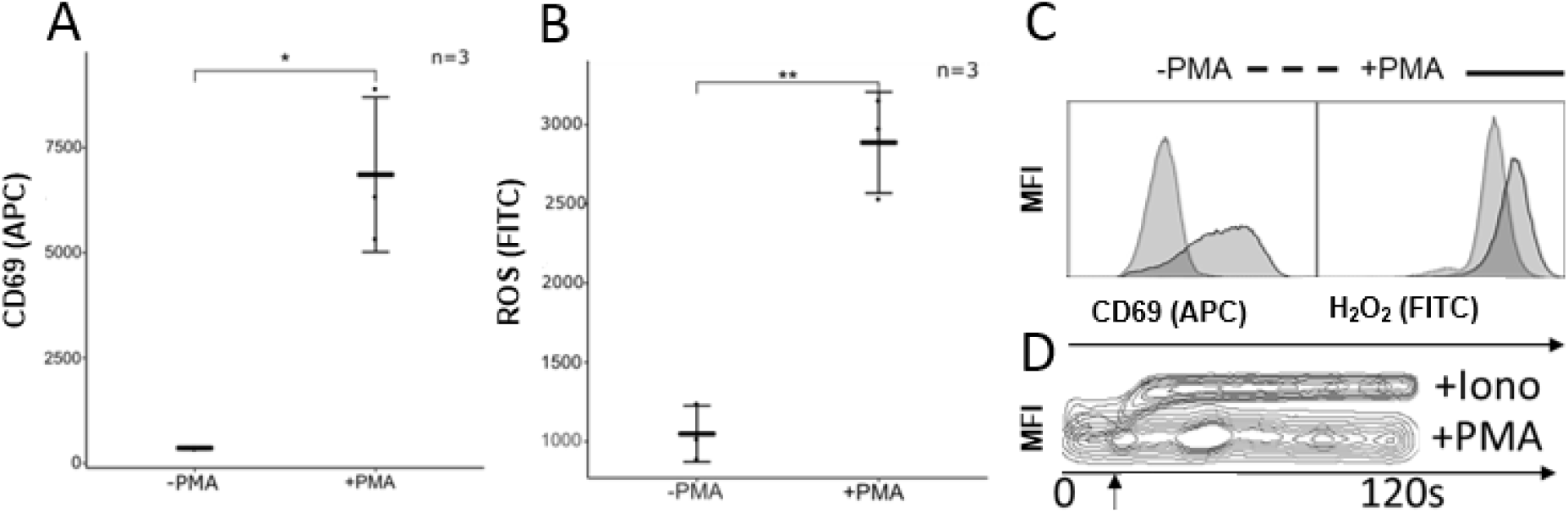
PMA stimulation induced mtROS signaling without inducing calcium signaling. Isolated CD3^+^CD4^+^ were activated with 10 ng/ml PMA for 1 hour (A-C) and analyzed for T cell activation as seen by CD69 surface-expression (A), and mtROS production as seen by H_2_DCFDA fluorescence (B). (C) Representative flow cytometry data showing CD69 surface expression (left panel) and mtROS production (right panel). (D) Intracellular calcium was measured using Fluo-4FF. Arrow indicates the application of PMA or Ionomycin. (A, B) Statistical analysis was performed using a two-sided, paired T-test, * p<0.05, ** p<0.01, n=3.

In parallel, mtROS production was examined in the same isolated CD3^+^CD4^+^ T cell samples. Following PMA stimulation as described before and using the ROS sensor H_2_DCFDA and flow cytometry, the mean fluorescence intensity (MFI) of the H_2_DCFDA signal increased from 1100 to 2750 confirming robust induction of ROS signaling in isolated T cells as well (Figure 1B). Interestingly, ROS signals exhibited notable low donor variation indicating physiological levels required for optimal immune responses. Since calcium signaling can impact ROS signaling (66) and thus might interfere with our mtROS dependent redox proteome approach, we carefully validated potential calcium responses in PMA-treated primary T cells. Free intracellular calcium was stained and quantified by flow cytometry using fluor-4FF, indeed confirming that activation of T cells with PMA does not yield any increase in intracellular calcium levels (Figure 1C).

Finally, we confirmed that stimulated primary T cells did not develop an apparent stress phenotype (Supplementary Figure 3A). Furthermore, we validated whether ROS emanates from mitochondria, and as expected, microscopy indeed confirmed that ROS induction was always co-localizing with the mitochondrial compartment (Supplementary Figure 3B). Finally, we utilized a catalase control assay to confirm that the majority of ROS produced by the T cell is H_2_O_2_ (Supplementary Figure 3C).

Thus, the applied stimulation conditions on of healthy individuals indicated CD3^+^CD4^+^ T cells as suitable to study thiol-specific redox modifications, which coincide with H_2_O_2_, by proteomics.

mtROS-generating CD4^+^ T cells maintain their overall thiol redox homeostasis To globally characterize reversible mtROS sensitive network components such as sulfenic acids (R-SOH) and disulfide bonds (R-S-S-R’), we applied a thiol switching method in which redox modifications of free/reduced as well as reversibly oxidized cysteine residues could be identified and quantified. In brief, free cysteine thiolates were conjugated with an iodoacetyl group coupled to an MS^2^-specific reporter mass (iodoTMT) which is liberated during peptide sequencing by mass spectrometry allowing comparative quantitation of oxidations. For the redox proteome approach in this study, CD4^+^ T cells were isolated in high purity from the PBMC fractions of healthy donors and samples of each donor were stimulated for 1 hour with 10 ng/ml PMA or left unstimulated (Figure 2A; step 1). In the first step of the thiol switching method, T cells were chemically lysed in the presence of different iodoTMT reporter tags allowing the direct labeling of free cysteine thiols (R-S^-^) in proteomes derived from unstimulated (CysFree^I^) or stimulated (CysFree^A^) T cells (Figure 2A; step 2). Subsequently, all samples were then reduced with tris(2-carboxyethyl)phosphine (TCEP) to yield free cysteine thiolate residues from previously reversibly oxidized sulfenic acids or disulfide bonds. Those now free thiolates were then subjected to a second round of labeling, again using iodoTMT reporter tags containing 2 different mass reporters for unstimulated (CysOx^I^) and stimulated (CysOx^A^) conditions, respectively (Figure 2A; step 3).

**Figure 2:**
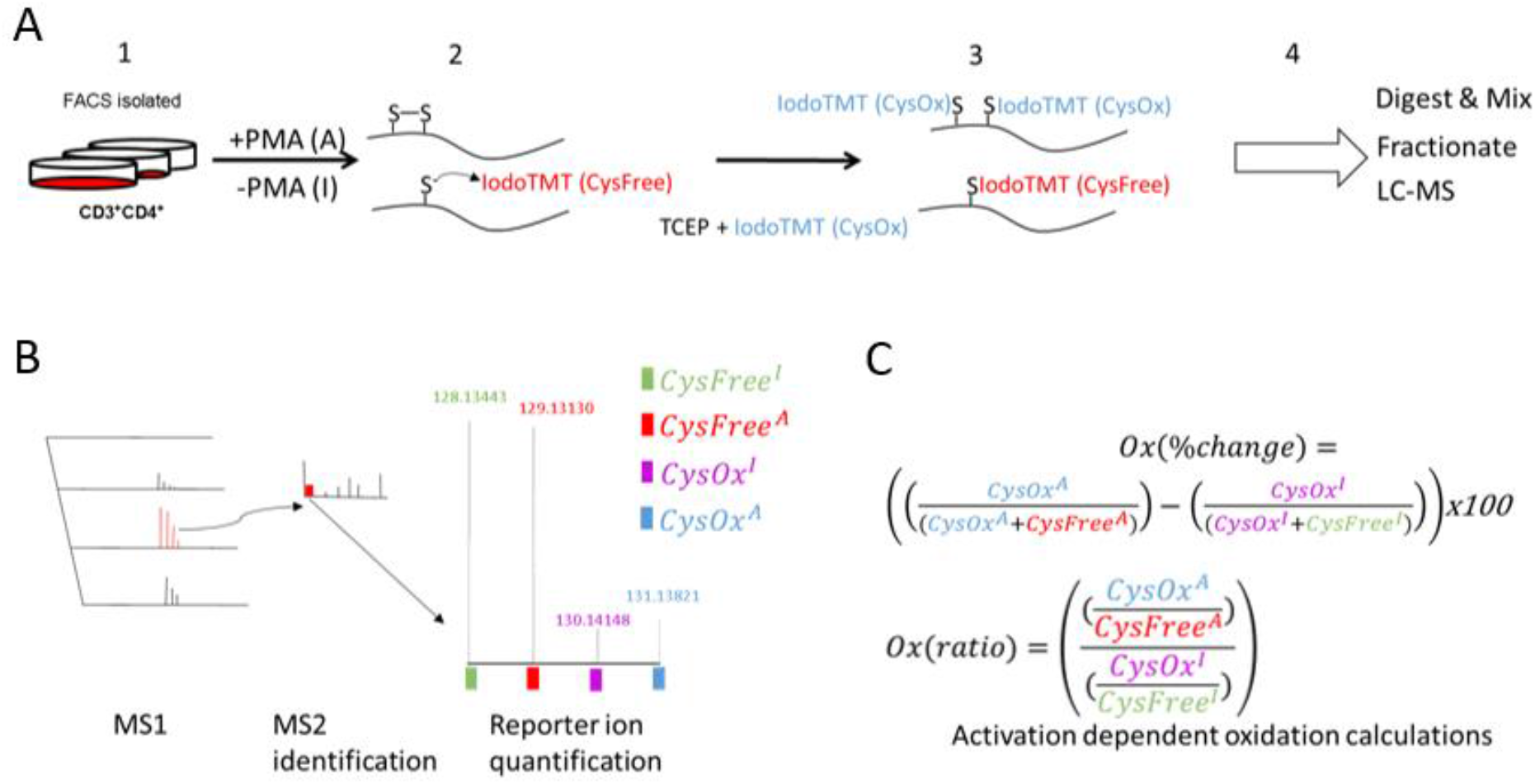
Multiplexed thiol-specific proteomic workflow that allows the quantification of residue specific cysteine oxidations. (A) CD3^+^CD4^+^ T cells were isolated from human PBMC by FACS-based cell sorting (1) and were either activated with 10 ng/ml PMA for 1 hour (A) or left inactive (I). Cells were lysed in the presence of a thiol reactive (iodoTMT) tag, which labeled any free cysteine thiolate residues (2). After the removal of excess label, samples were reduced by TCEP resulting in the conversion of all reversibly oxidized cysteine residues (such as disulfide bonds or sulfenic acids) to free cysteine thiolate residues, which were subsequently labeled with another iodoTMT tag with a differing reporter mass (3). Lysates were mixed and digested before being subjected to offline high pH reverse phase chromatography to generate multiple fractions, which were then analyzed under conventional low pH reverse phase nanoLC-MS (4). For simplicity, (2 and 3) only shows the iodoTMT tag strategy for the activated (A) sample. (B) MS1 peptide mass fingerprints were taken for subsequent activation experiments utilizing Higher-energy C-trap dissociation (HCD) fragmentation to generate high quality MS^2^ spectra. By the use of the reporter tags that were identified in the MS^2^, cysteine-containing peptides were identified and quantified using automated software, and quantitative information for 9598 cysteine-containing peptides could be generated from 3284 proteins. (C) Formulas used for the calculation of (i) the total % change in oxidation of each residue between stimulated versus unstimulated conditions or (ii) the fold change in oxidation between stimulated over unstimulated conditions.

In order to compare the thiol redox proteome in these discrete conditions, differentially iodoTMT-labeled protein samples were combined donor-dependently and subjected to tryptic digestion and high pH reverse phase fractionation to improve proteome coverage (Figure 2A; step 4). Identification and quantification of cysteine modifications were then accomplished by peptide ion activation experiments wherein both parent and fragmentation ions were characterized by high-resolution MS (Figure 2B, Supplementary Figure 4). In total, shotgun proteomics identified 3284 proteins from which 9598 cysteine-containing peptides were associated, of which 4784 cysteine-containing peptides could be quantified in at least 2 of 3 donors under all conditions (Supplementary Table 1). Fragmentation spectra of each donor unambiguously identified the sequence of cysteine-containing peptides and comprised four reporter ion intensities indicating the amount of free or oxidized cysteine residues in nonstimulated or stimulated conditions.

The robustness of our experimental approach was determined by looking on the sample specific IodoTMT tag distributions. Intensities of reporter ions derived from tags used to label free cysteine residues (CysFree) were found consistently more abundant than those from tags used to label reversibly oxidized thiols (CysOx). Generally, there was only moderate inter-sample deviation of the same conditions between different donors (Supplementary Figure 5). It is well known that the majority of cysteine residues in proteins exist as either significantly oxidized or significantly reduced (67). The oxidation state reflects the overall thiol redox homeostasis of the compartment, with cytosolic proteins having mostly reduced cysteine residues, and conversely, extracellular proteins typically contain structural disulfide bonds. Only a few exceptions exist: redox active thiol residues, typically found in redox active antioxidants such as thioredoxins or peroxiredoxins, can be found in intermediate oxidation states, reflecting their catalytic importance to maintain redox homeostasis. To validate this, we generated a histogram view of the average cysteine oxidation states of the thiol residues derived from our T cell proteomes. Indeed, our data was found consistent with previous literature and displayed a valley shape distribution with a large number of cysteine residues at the extreme ends (<30% oxidized and >70% oxidized, Figure 3).

**Figure 3:**
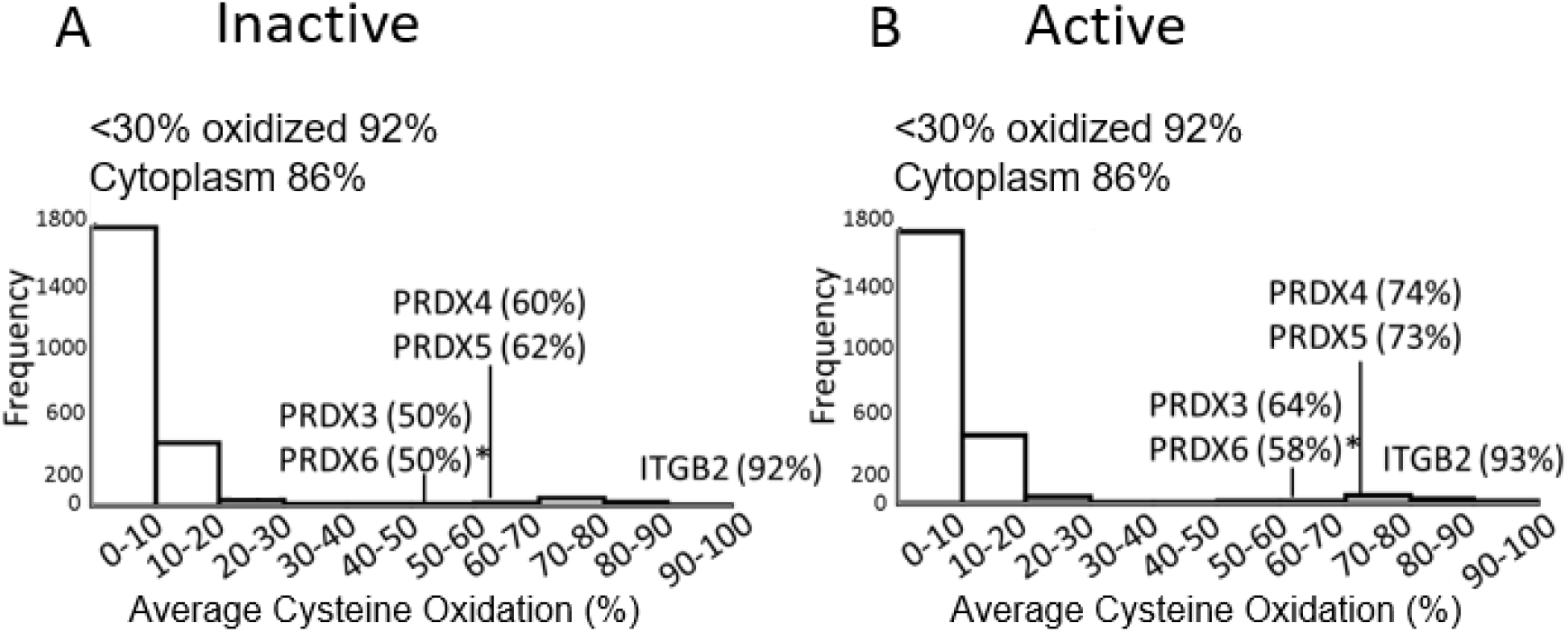
Distribution of average oxidation of cysteine residue-containing peptides in (A) unstimulated and (B) stimulated CD4+ T cells. PBMCs were isolated from healthy donors and stimulated with 10 ng/ml PMA for 1hour. Average oxidation of cysteine residues was calculated according the formula given in Figure 2C, which is ((CysOx^l^/ CysOx^l^+CysFree^l^)*100) for unstimulated (inactive) cells (A) and ((CysOx^A^/CysOx^A^+CysFree^A^)*100 for stimulated (active) cells (B). Selected candidate proteins found with significantly differentially oxidized cysteine residues between stimulated and unstimulated conditions are marked with an asterisk. Statistical analysis was performed using a two-sided, paired T-test, * p <0.05, n=3.

We next compared the average oxidation state of sensitive thiols in both unstimulated and PMA stimulated conditions (Figure 2C, upper equation). Subsequently, we used iodoTMT reporter ion intensities to calculate the average percentage change of oxidation for each peptide. The overall distribution of the thiol oxidation states under both conditions was found notably similar and in accordance with results from other cell types (21,67). In both the stimulated and unstimulated CD4^+^ T cells, 92% of the characterized cysteine residues were found predominantly reduced with an oxidation status of less than 30%, with the majority residing in the 0-10% oxidation range (Figures 3A & 3B). The majority (86%) of reduced cysteine residues reside in proteins that are localized in the cytoplasm according to the panther gene ontology database (68,69). Whereas, non-cytoplasmic cysteine residues in the <30% range were mostly found in proteins that are annotated as components of the mitochondrial matrix (7%), and only a few cysteine residues (0.8%) could be associated to proteins localizing at the highly oxidizing endoplasmic reticulum (70). Conversely, highly oxidized cysteine residues with an oxidation status higher than 90% were mostly found in surface localized proteins containing structural disulfide bonds such as integrin beta chain-2 (ITGB2/CD18) (Figure 3).

Thus, the overall oxidation confirmed CD4^+^ T cells as able to maintain global thiol redox homeostasis upon stimulation. Indeed, several antioxidant proteins were identified by our approach including thioredoxin (TXN), thioredoxin-like proteins (TLXN) and peroxiredoxins (PRDX) which exhibited intermediate oxidation states (Supplementary Table 1). Of the total characterized cysteine-containing peptides 18 cysteine residues were found significantly differentiated (Supplementary Table 2). We manually validated which of those underwent a shift in oxidation in our mtROS producing T cells suggesting their role in preserving T cell redox homeostasis. This revealed a shift of oxidation particularly at peroxiredoxins, that are known to play a key role in redox dependent signaling (53). Peroxiredoxins are thiol-specific peroxidases and utilize conserved redox sensitive active sites consisting of one active cysteine sulfenic acid (PRDX1-4 and PRDX6) (71,72) or a cysteine disulfide bond (PRDX5) (73) used during the catalytic conversion of H_2_O_2_ to H_2_O. Indeed, peroxiredoxins (PRDX3/4/5/6) identified in our study were found to increase oxidation by an average of 10% at catalytic cysteine residues following T cell activation (Figure 3A and B). Of those, the shift of oxidation was found mostly conserved and significant for PRDX6, which exhibited an increase in oxidation at its active site cysteine (Cys47) of 8%.

Thus, the average oxidation profile of CD4^+^ T cells was found prototypic to other cell types and confirmed their ability to maintain global thiol redox homeostasis upon stimulation and mtROS production. Antioxidative proteins and redox active cysteines were identified with intermediate oxidation states and peroxiredoxins, in particular PRXD6, seem to play a major role contributing to redox homeostasis.

### Identification of mtROS-sensitive processes

Following classification of the global average cysteine oxidation in the CD4^+^ T cell populations, we then aimed to identify redox-sensitive cysteine thiolates, which are most differentially oxidized following the generation of an mtROS signal from the mitochondria. To this end, the degree of oxidation for each cysteine thiol was comparatively quantified in stimulated and unstimulated CD4^+^ T cells. Regulations were determined utilizing the fold changes of the ratio of the oxidized cysteine (CysOx) over the reduced cysteine (CysFree) in a given peptide upon T-cell stimulation (see Figure 2C, lower equation). The majority of oxidized cysteine residues in individual donors were detected with a difference between −0.5 and 0.5 log2. Noticeably, there was a higher induction of stimulation induced oxidations as seen by the shift towards positive regulatory values. As such, we considered cysteine residues as H_2_O_2_-sensitive that were found consistently regulated in at least 2 of 3 donors with a p-value less than 0.05 and a log2 fold change greater than 0.5, similar as used in other studies (74). Our study revealed 88 differentially oxidized cysteine residues at 76 cysteine-containing peptides, as certain peptides contained more than 1 modified cysteine residue. Ultimately, all cysteine-containing peptides were associated to 71 proteins as certain proteins contained multiple significant peptides in PMA-stimulated T cell samples as compared to unstimulated T cell samples (Figure 4, Supplementary Table 3). Interestingly, both oxidation and reduction of cysteine residues were detected, but oxidations were found more often and with higher p-values. Specifically, 70 cysteine residues from 62 cysteine-containing peptides were characterized significantly more oxidized following T cell stimulation, whereas only 18 cysteine residues from 14 cysteine-containing peptides demonstrated a shift towards a more reduced state (Figure 4).

**Figure 4:**
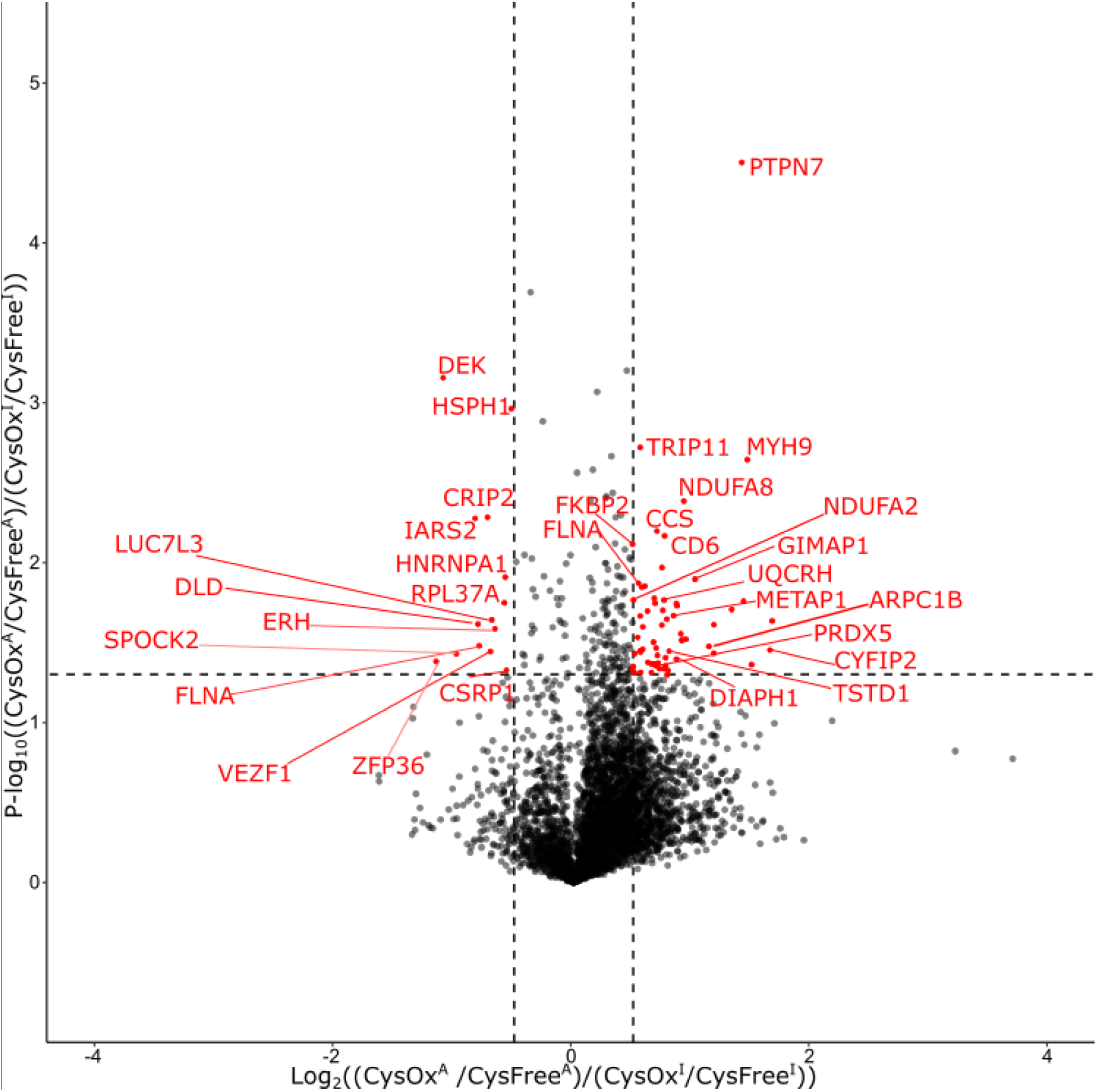
Regulation of redox-modified CD4^+^ T cell proteins following PMA stimulation. Volcano plot showing proteins with regulated reduced (left) and oxidized (right) cysteine residues upon PMA stimulation of CD4^+^ T cells. PBMCs were isolated from healthy donors and stimulated with 10 ng/ml PMA for 1 hour, fold changes (Oxratio) in cysteine oxidation were analyzed as depicted in Figure 2C. In total, 4784 cysteine-containing peptides could be quantitatively analyzed of which 76 cysteine-containing peptides originating from 71 proteins were found either differentially oxidized (62) or differentially reduced (14) upon activation (highlighted in red) in at least 2 of 3 donors. X-axis shows the fold change (log_2_) for each cysteine residue, y-axis shows the p-value (−log10) for each cysteine residue. The cut-off for fold change was set at log_2_ 0.5, and statistical significance was determined with a two-sided, paired t-test, and a p-value of −log10 1.3 (p< 0.05) was taken as significant, n=3.

Using these 71 regulated proteins, we then performed enrichment analyses to evaluate which processes and pathways mostly undergo oxidation or reduction at their components. String database (59) analyses revealed the majority of oxidized proteins involved in cytoskeleton dynamics and immune functions (Figure 5A) indicating these networks to be regulated by mtROS. In total, protein network analyses and the inspection of assigned cellular functions allowed to group proteins with induced oxidations in seven functional categories. Besides cytoskeleton dynamics and immune functions, oxidized proteins were found to be involved in RNA processing, function as antioxidants (such as PRDX5) or mediate post-translational modifications, whereas only 9 oxidized proteins belong to mitochondrial functions or support ER processing.

**Figure 5:**
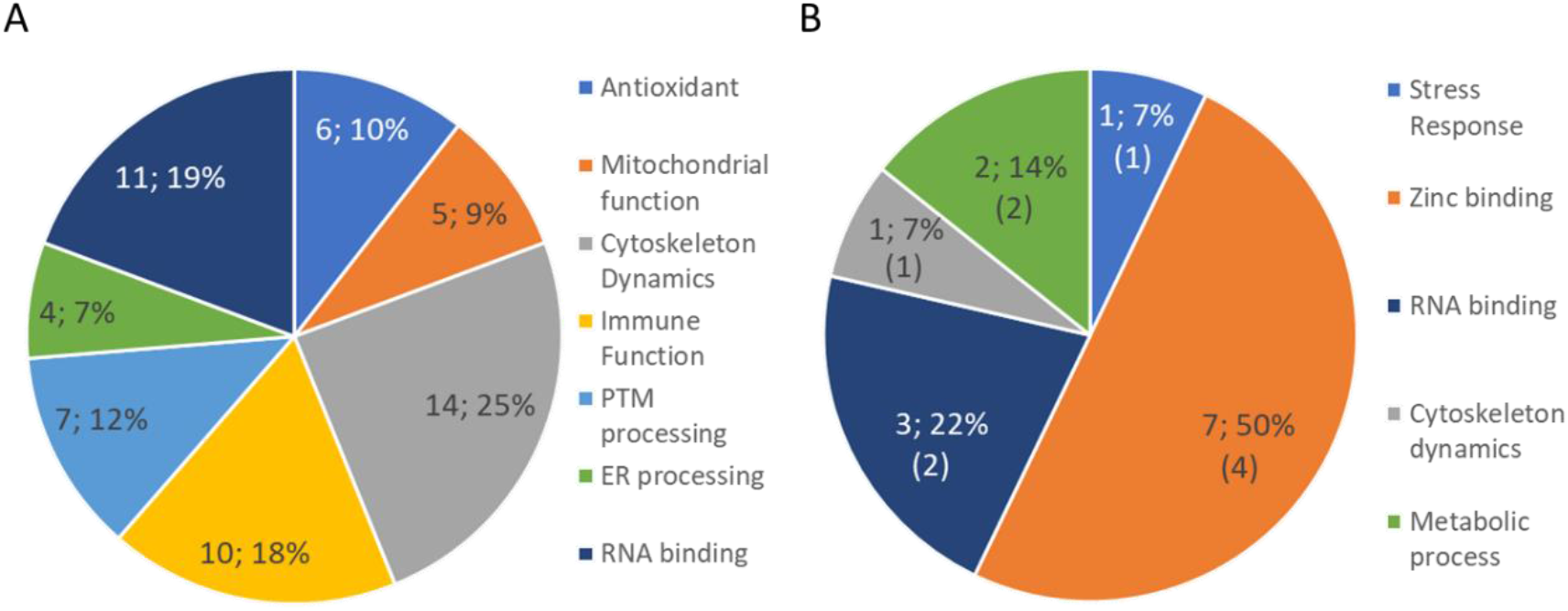
Cellular processes modified by oxidation or reduction in mtROS-producing CD4^+^ T cells. Enrichment analyses considered the 71 proteins with differentially redox-modified cysteine residues (from Figure 4) and revealed cellular processes mostly affected by oxidation (A) or reduction (B). The class “immune function” is presented for oxidized proteins only, whereas the majority of reduced proteins were related to immunity before. As such, the left most number indicates the actual value of proteins per category while the number of immune related proteins in (b) indicate the number of immune related proteins per category.

Conversely, five functional processes could be defined in the group of reduced cysteine residues. Additionally to the categories mentioned before (antioxidants, RNA processing and cytoskeleton dynamics), reduced proteins were found to play a role in metabolic processes or exhibited zinc-binding motifs. Zinc-binding proteins were found to be the most enriched functional category (50%, Figure 5B). It was demonstrated that zinc-finger cysteine residues exist as thiolates similar to other redox active proteins. Zinc-fingers are therefore known to be susceptible to oxidation by H_2_O_2_ (75). Of interest, most of the affected proteins were related to immune functions before, and thus might function as novel thiol switch proteins for immune modulation.

Next, we evaluated which components in the group of 71 proteins underwent the most robust regulations with a minimum of donor-variation (highest p-value, Supplementary Table 3). When focusing on the top 10 robustly regulated oxidized peptides (Table 1), four were found to be cytoskeleton or cytoskeleton-associated proteins, including proteins which associate with the wave regulatory complex (ARPC1B, FLNA, DIAPH1 and CYFIP2) as well as the myosin complex (MYH9). Notably, CYFIP2 is the T cell specific variant (76), demonstrating the specificity of the analysis.

**Table 1:** Top 10 most robustly differentially oxidized (Ox(ratio)) proteins. Listed are the identified sequence (modified cysteine residues in red), gene name, fold change (log2), percent change, as well as major protein function and localization. Proteins are listed in order of significance (see Figure 4 and Supplementary Table 3).

| Sequence/Position | Gene name | Fold change (Log<sub>2</sub>) in oxidation | % Change in oxidation | Function | Localization |
| --- | --- | --- | --- | --- | --- |
| SLGAVEPIC<sub>62</sub>SVNTPR | PTPN7 | 1.42 | 10 | Immune response (80) | Cytoplasm |
| LVLAC<sub>686</sub>EDVR | TRIP11 | 0.57 | 5 | Cytoskeleton dynamics (81) | Golgi |
| LQLQEQLQAETE<br>LC<sub>896</sub>AEAEELR | MYH9 | 1.47 | 1 | Cytoskeleton dynamics (82) | Cytoskeleton |
| AAAHHYGAQC<sub>36</sub>DK<br>PNK | NDUFA8 | 0.93 | 8 | Oxidative phosphorylation (78) | Mitochondrial inner membrane |
| QIC<sub>244</sub>SC<sub>246</sub>DGLTIWEE<br>R | CCS | 0.71 | 4 | Antioxidant (51) | Cytoplasm |
| LC<sub>137</sub>EVVEHAC<sub>144</sub>R | CD6 | 0.77 | 5 | Immune response (83) | Cell membrane |
| GWDQGLLGMC<sub>97</sub>E<br>GEKR | FKBP2 | 0.50 | 7 | ER Processing (84) | ER |
| SHTEEDC<sub>67</sub>TEELFD<br>FLHAR | UQCRH | 0.70 | 7 | Oxidative phosphorylation (85) | Mitochondrial inner membrane |
| C<sub>76</sub>HVEVVDTPDIFSS<br>QVSK | GIMAP1 | 1.03 | 10 | Immune Response (86) | Endoplasmic Reticulum |
| C<sub>444</sub>SYQPTMEGVHTV<br>HVTFAGVPIPR | FLNA | 1.55 | 5 | Cytoskeleton dynamics (87) | Cytoskeleton |

However, the most robustly regulated cysteine oxidation was identified at the protein tyrosine phosphatase 7 (PTPN7), which interacts with MAPK1 and additionally regulates T- and B-lymphocyte development (77). Additionally, respirasome components were oxidized with lowest donor-variations, these include the NADH dehydrogenase [ubiquinone] 1 alpha subcomplex subunit 8 (NDUFA8) and the ubiquinol-cytochrome c reductase hinge protein (UQCRH) which both are known regulators of electron transport at complex I and III, respectively (78,79).

Conversely, when focusing on the top 10 most robustly reduced cysteine-containing peptides (Table 2), the class of zinc-binding proteins was, in a similar manner to the unbiased approach (Figure 5B), enriched (6/10). Of these, ZFP36 and CRIP2 have already been implicated to impact T cell activation (88,89) through transcriptional regulation, whereas the others might be new candidate proteins that may regulate full T cell activation. Zinc ions are known to stabilize the chelating thiolate and yield a mtROS sensitive thiolate residue (75). Thus, our proteome data indicated that H_2_O_2_ oxidized the coordinating cysteine thiolates in the zinc-finger motif resulting in the zinc-exclusion prior to their subsequent reduction by anti-oxidative T cell responses or antioxidative environments (see discussion). Four out of the 10 top regulated reduced proteins were annotated to be nuclear localized, a subcellular region known to be resistant to oxidation (90). Interestingly, the mitochondrial proteins identified in the group of top reduced candidates were IARS2 and DLD, both of them residing in the reducing compartment of the mitochondrial matrix (91). IARS2 is responsible for synthesizing mitochondrial tRNA and has been reported to impact steady state abundances of various ETC protein subunits (92). DLD is a homodimer known to mediate a number of metabolic processes in the mitochondria including the oxidation of dihydrolipoamide (93).

**Table 2:**
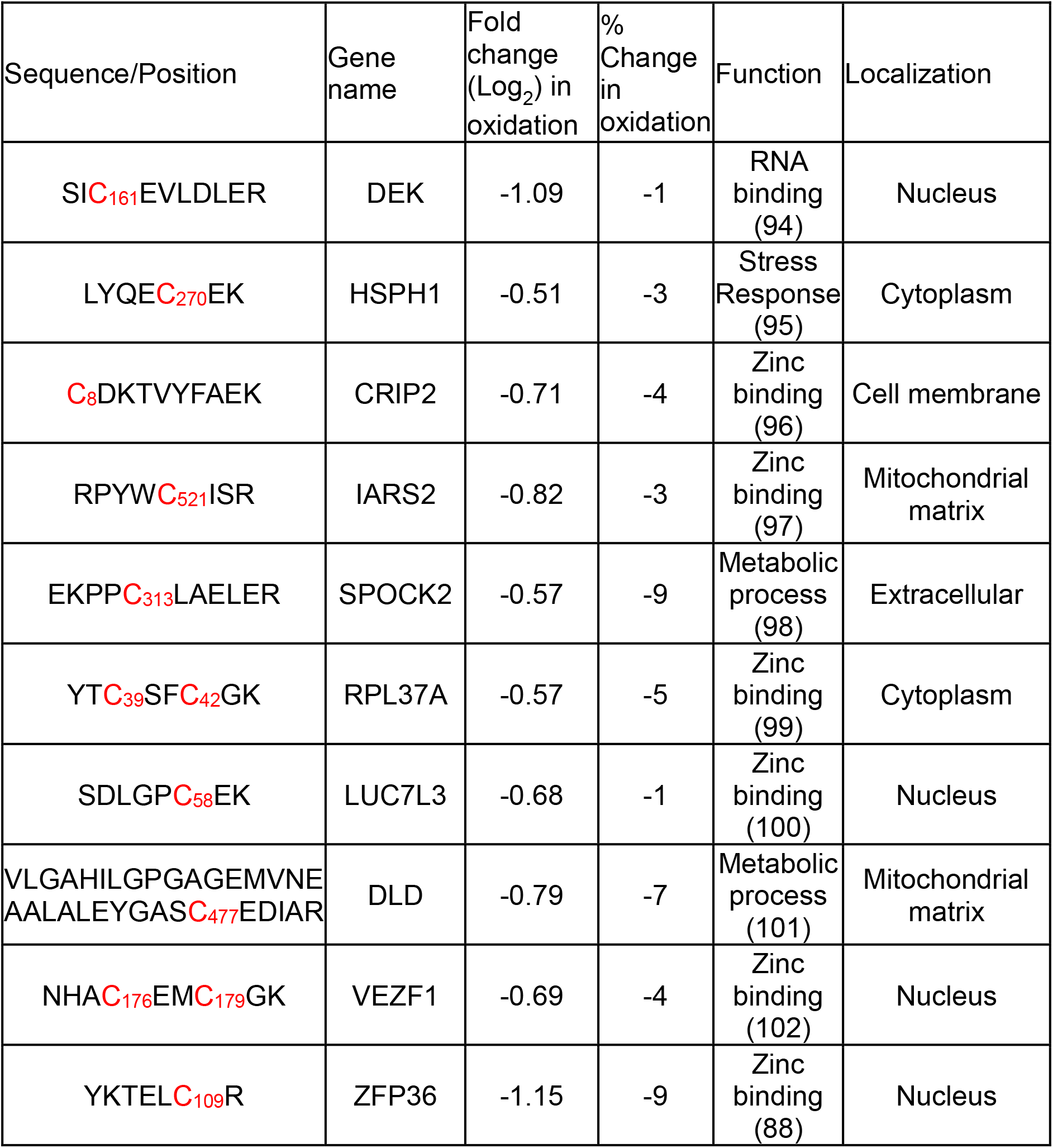
Top 10 most statistically significant differentially reduced (Ox(ratio)) proteins. Listed are the identified sequence (modified cysteine residues in red), gene name, fold change (log2), percent change, as well as major protein function and localization. Proteins are listed in order of significance (see Figure 4 and Supplementary Table 3).

As the mtROS in T cell activation seemed to emanate from the mitochondria (see Supplementary Figure 3B), the microenvironment surrounding the mitochondria could be considered as oxidizing. And since zinc-finger proteins constituted the most enriched reduced candidates, we asked the question whether H_2_O_2_ induce zinc exclusion from the zinc-fingers in stimulated T cells, and whether we can detect free zinc near to mitochondrial microenvironments. To answer this question, we isolated PBMCs and stimulated T cells with 10 ng/ml PMA in the presence of the cell-permeable fluorogenic zinc reporter Zinpyr-1 and the mitochondrial dye MitoTracker red. Cells were prepared for immunofluorescence microscopy, and CD3^+^ T cells were imaged for distribution of free zinc and mitochondria. Stimulated T cells as part of PBMC fractions could be unambiguously characterized by the expression of CD3 at the surface (data not shown) and, as expected, T cells did not indicate polarization of CD3 or mitochondria. When considering cross-sectional intensities of mitochondria, we observed that free zinc almost but not exclusively co-localized with the mitochondrial microenvironment (Figure 6 and Supplementary Figure 6).

**Figure 6:**
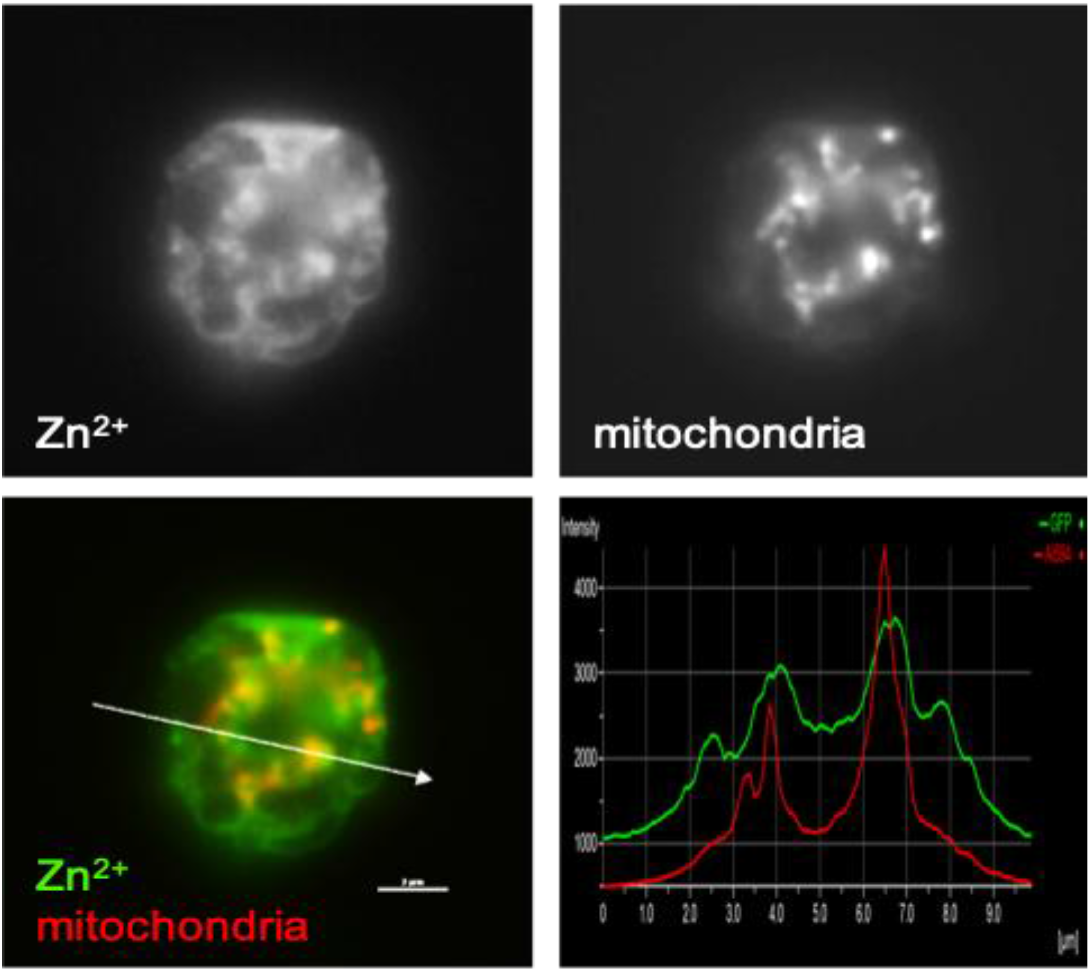
Localization of free zinc ions (Zn^2+^) and mitochondria in mtROS-producing T cells. PBMCs were stimulated with 10 ng/ml PMA for 1 hour in the presence of Zinpyr-1 and MitoTracker, prepared for fluorescence microscopy, and cross-sectional analyses of fluorescent signals were analyzed in CD3^+^ T cells (n=40). Representative images showing the local distribution of free Zn^2+^ (top left panel, and green in merged image) and spatial clustering of mitochondria (top right panel, and red in merged image). Fluorescence intensity plots revealed local co-emerging of free zinc at the mitochondria (lower right panel). Arrow indicates the plane of the intensity plots, scale bar = 2μm.

To summarize, we identified a repository of candidate T cell-specific mtROS-sensitive thiols associated to both oxidized and reduced proteins. In particular, the mitochondrial respirasome, actin cytoskeleton and proteins related to immune functions were found to be mtROS sensitive and appeared highly oxidized upon T cell stimulation. Conversely, zinc-finger proteins and mitochondrial matrix associated proteins were enriched in the candidate proteins that were reduced after stimulation. Reduction of identified zinc-coordinating cysteine residues strongly suggested zinc exclusion and microscopy confirmed free zinc ions enriched at the mitochondrial microenvironment of mtROS-producing T cells, implicating mtROS as a potential transcriptional regulator.

## Discussion

T cells are professional ROS (reactive oxygen species) producing cells which have been demonstrated to utilize the mitochondria as a signaling organelle and require mitochondrial-derived ROS (mtROS), in particular H_2_O_2_, to fulfill their physiological immune responses (103). H_2_O_2_ is uniquely reactive to functional cysteine thiolates that, in T cells apparently affects different immune pathways.

H_2_O_2_ has been found to modulate non-redundant functions in different T cell subsets including the activation (104) and cytotoxic function (105) of CD8^+^ T cells as well as the differentiation of CD4^+^ Treg cells (106). We found conventional CD4^+^ T cells of healthy individual’s demonstrated superior mtROS inducibility as compared to CD8^+^ cytotoxic T cells, concurring with subset specific mtROS function. In general, H_2_O_2_ is nowadays well accepted to act as a second messenger in immunity. The best characterized T cell specific function being NFAT signaling (7,41,107,108), which has triggered research focusing on mtROS sensing mechanisms (5,12,109) including H_2_O_2_ sensitive thiol switches.

In this line, results of this study now complement our understanding on the role of mitochondrially produced H_2_O_2_ in adaptive T cell immunity. H_2_O_2_ production did not cause a global shift of the proteome redox state in T cells but in fact coincided with discrete oxidations of a number of molecular functions including the mitochondria, actin cytoskeleton and zinc-binding in particular.

### Fine-tuning mtROS along metabolic reprogramming of T cells

This study identified a number of mtROS-sensitive processes which suggest novel regulatory mechanisms in activated T cells. Among those, we complement information of electron leakage sites at the respirasome and identified network components that indicate how activated T cells can fine-tune mtROS generation in and outside the mitochondria.

Electrons can leak out of both complex I (6,110,111) and complex III (8,112,113) of the respirasome, which could indeed be supported by our data. At complex I we found NADH dehydrogenase [ubiquinone] 1 alpha subcomplex subunit 2 (NDUFA2) and NADH dehydrogenase [ubiquinone] 1 alpha subcomplex subunit 8 (NDUFA8) to be significantly oxidized (+2% and +9% shifts at Cys58 and Cys36, respectively) in PMA stimulated T cells. NDUFA2 is a component of the CI electron transfer arm between the NADH electron donator and the coenzyme Q site (114). NDUFA8 is a part of the intermembrane arm which serves to pump out protons into the intermembrane space (115), and additionally, Cys36 of NDUFA8 has been shown to contain redox sensitive thiols (116). Both components are in close proximity to the Coenzyme Q binding site which has been implicated in leaking electrons (117), we now hypothesize that they leak out of CI near the sites where NDUFA2 and NDUFA8 localize leading to their oxidation.

With respect to Complex III, Cytochrome c1 (CYC1) has been implicated as a site of electron leakage and participates in electron transfer reactions with cytochrome c (9,118,119). Interestingly, the metabolic protein Dihydrolipoyl dehydrogenase (DLD) was reported to be uniquely reduced by mtROS that emanates from CIII (120). We observed DLD to undergo modification following PMA stimulation at Cys477 (−7%).

This indicated a novel redox active site of DLD which may mediate its activity since this cysteine is on the interface of the interaction site of the homodimer (121).

We theorize that the electrons which leak from complexes I and III mediate an intra-mitochondrial fine-tuning mechanism, primarily through conformational changes in CIII. Indeed, modulation of CIII structure to alter respirasome function has been shown by a number of different groups (122–124). Upon PMA stimulation, we observed a strong oxidation at Cys53 (+8%) of the structural disulfide of the complex III subunit, ubiquinol-cytochrome c reductase hinge protein (UQCRH) (85,125,126). UQCRH is directly associated to CYC1 in Complex III and has been shown to mediate a stronger interaction between Cytochrome c and CYC1 through the unique sequence of 8 glutamate residues near its C-terminus (125–128). Cys53 of UQCRH in particular was demonstrated by others in bovine heart mitochondria to be constitutively oxidized to structural disulfides with Cys67 (127). This disulfide in UQCRH mediates a favorable position of the glutamic acid-rich sequence near the transfer active site of CYC1 and Cytochrome c for efficient electron transfer and subsequently diminished electron leakage (126). This is exemplified by the reduced cellular respiration in the absence of UQCRH although the structure of the super complex is maintained (129), suggesting it may not be critical for cellular respiration, however, may in fact be a respiration regulator with a mtROS-sensitive thiol switch. Indeed, it has been shown that under periods of oxidative stress, CIII and more specifically UQCRH are particularly regulated by oxidation (79), a situation similar in T cell activation.

There is growing evidence on the importance of the cellular microenvironment for regulating the respirasome. Here, our data suggests cytoskeleton components to potentially establish extra-mitochondrial feedback systems to regulate respiration and mtROS production. For instance, actin cycling involving the Actin-Related Proteins (ARP2/3) complex regulates respirasome dynamics by fission or fusion of mitochondria (130). Interestingly, we identified the ARP2/3 complex subunit ARPC1B (Supplementary table 3) to be oxidized at its residues Cys342 and Cys346. ARPC1B expression is restricted to hematopoietic cells and germline mutations of ARPC1B recently were demonstrated to affect T cells causing immunodeficiency (131,132). Data of our study suggest mtROS as a mechanistic link between how T cells involve the ARP2/3 complex in activation and implicate Cys342 and Cys346 of ARPC1B (+2%) as novel H_2_O_2_ sensitive thiols which may potentially mediate interactions with several candidate proteins, including Cytoplasmic FMR1-interacting protein 2 (CYFIP2, Supplementary table 3) that was found oxidized at positions Cys1087 and Cys1088 (+7%). CYFIP2 interacts with the ARP2/3 complex alongside Ras-related C3 botulinum toxin substrate 1 (RAC1), which is an activator of the Wave Regulatory Complex (WRC) (133) and transduces signaling to ARP2/3, suggesting a thiol switch. Additionally, Filamin A acts as an essential scaffold with the ARP2/3 complex to mediate actin branching (87) and was found oxidized at two distinct peptides, both Cys444 and Cys2160 (+5% and +15%, respectively) highlighting potentially multiple sites of interaction between ARP2/3 and FLNA. Additionally, Protein diaphanous homolog 1 (DIAPH1) was found oxidized at Cys1227 (+7%), which also can mediate mitochondrial trafficking (134), as well as immunological synapse formation in T cells with ARP2/3 (135) showing the impact in T cell function.

Additionally, subunits of the cytoplasmic Condensin II complex were reported to impact respiration and oxidative stress responses in mitochondria whereby individual subunits can be imported to mitochondria (136,137). Our experiments identified the Condensin II complex subunit Bifunctional aminoacyl-tRNA synthetase (EPRS), however, the regulatory information obtained from one donor sample was missing. Although EPRS might be a potential mtROS sensor that mediates a negative feedback loop, its oxidation at Cys1480 (+5%) was not found statistically significant in this study (Supplementary table 1).

Furthermore, on a global level, we did not observe a global shift in the T cell redox state upon PMA activation. This could be supported by the shift in oxidation of the peroxiredoxin 5 (PRDX5, Cys204, +10%) and peroxiredoxin 6 (PRDX6, Cys47, +8%) which were oxidized at their catalytic cysteine residues. Peroxiredoxins are the preferential targets of H_2_O_2_ (53) and peroxiredoxins such as PRDX5 and PRDX6 are known to primarily catalyze the conversion of H_2_O_2_ to H_2_O in the mitochondria (138) or in other compartments such as the cytoplasm (139). In parallel to their antioxidant function, peroxiredoxins mediate signaling events through the reversible oxidation of thiol switch proteins, including phosphorylation events through inhibition of protein tyrosine phosphatase proteins (140–142). Additionally, other phosphorylation signaling cascades via the apoptosis signaling kinase1 (ASK1) and mitogen activated protein kinase (MAPK) can be modulated by peroxiredoxins (143,144). We hypothesize that during T cell activation PRDX 5 and 6 are oxidized to maintain global redox homeostasis in CD4^+^ T cells while mediating additional redox signaling affecting phosphorylation signaling in T cells to facilitate immune function.

It was interesting to note that antioxidants seemed to undergo more regulatory changes than large absolute shifts in oxidation, which was evident by the number of antioxidant proteins identified utilizing the Ox(ratio) equation (Figure 2C, lower equation). Alongside PRDX5, a few components involved in glutathione activity have been identified including Thiosulfate:glutathione Sulfurtransferase (TSTD1), a reducing agent of both glutathione (GSH) and thioredoxin (TXN) (145). TSTD1 was found oxidized at Cys79 (+3%) that is its functional and sensitive thiol. Also impacting glutathione is Methionine aminopeptidase 1 (METAP1) which has been demonstrated to control glutathione redox state (146) and was found oxidized at Cys179 (+5%). Additionally, Copper chaperone for superoxide dismutase (CCS) which incorporates the catalytic copper cation into the Superoxide dismutase [Cu-Zn] (SOD1) was found oxidized at its catalytic Cys244 (+4%). This observation suggests that multiple processes are necessary to ensure a global redox homeostasis.

### Zinc-finger motifs as putative mtROS dependent transcription regulators

Parallel to oxidation sensitive thiols, our iodoTMT approach identified a number of “reduced” cysteine residues. Cysteine residues within a zinc-finger motif are susceptible to oxidation due to the conjugated metal stabilizing the coordinating cysteine thiolate (22,147,148). Following the increase in oxidation, antioxidant proteins have been demonstrated to reduce zinc binding sites (149). As has been described for HSP33 (23,24,150), we hypothesize that the oxidation of the zinc-conjugating cysteine residues leads to exclusion of the free zinc, leading to differential activity of proteins during periods of high oxidative stress (150). Indeed, we showed that cysteine oxidation within zinc-binding proteins coincides with free zinc-rich clouds at the mitochondrial microenvironment. Thus, at the 1 hour time point in stimulated T cells zinc-binding cysteines have already released zinc and were subsequently reduced.

Here, studying primary CD4^+^ T cells with redox proteomics, we now identified proteins containing zinc-coordinating cysteine residues at the most enriched processes with respect to the reduced proteins (Table 2 and Supplementary Table 3). This observation thus strongly suggests zinc-binding as a novel mechanism of how oxidation states functionally can impact T cell immune function through transcriptional regulation as zinc-binding motifs such as the zinc-finger are known to bind DNA (151). It has been demonstrated before that under oxidative conditions generated by T cell activation, metallothionein proteins release their chelated zinc to allow for zinc as a signaling molecule (31). It is now well-accepted that zinc is an important component of T cell activation (29). For instance, following T cell stimulation, a local but specific zinc influx at the sub-synaptic space induces the binding of the lymphocyte-specific protein tyrosine kinase (Lck) to the CD4 and CD8 co-receptors in a so-called “zinc-clasp” motif-specific manner (30). Additionally, it has been demonstrated that PMA is capable of releasing free zinc into the cytoplasm in multiple immune cell subsets and zinc signaling was found critical for MAPK and NF-κB signaling (152).

Irrespective of the free zinc concentration, it is tempting to speculate that zinc-finger proteins regulate immune responses either by loss of transcriptional regulatory function or variant molecular interactions following zinc release. Indeed, a number of zinc-finger proteins underwent a change of oxidation at their redox sensitive thiol-containing cysteine residues (Supplementary Table 3). The mRNA decay activator protein 36 (ZFP36/ Tristetraprolin) is known to downregulate a number of T cell activator mRNAs including the CD4 receptor and Interleukin-2 (IL-2) required for clonal expansion in CD4^+^ T cells (153). Mechanistically, reduction of ZFP36 at Cys109 (-10%) was hypothesized to mediate upregulation of CD4^+^ T cell activation by loss of activity of the ZFP36 zinc-finger. As mentioned, ZFP36 can downregulate IL-2 activity (88), therefore inhibition of ZFP36 would upregulate CD4^+^ T cell colony stimulation, e.g. by stabilizing IL-2 transcription. Furthermore, the 60S ribosomal protein L37a (RPL37A) is known to impact the upregulation of cellular tumor antigen p53 (p53) through upregulation of the Mdm2-p53-MdmX network (154). p53 is known to downregulate CD4^+^ T cell activity in a TCR independent mechanism (155), similar to our PMA activation method. We speculate that interfering with the zinc-finger structure of RPL37A by oxidation of its Cys39 and Cys42 zinc-finger motif (−5%) could upregulate CD4^+^ T cell activity by downregulating transcription of p53. Additionally, upon T cell activation, Cysteine-rich protein 2 (CRIP2) can downregulate the activating transcription factor NF-κB activity by repressing its transcription (89). Therefore, loss of function of CRIP2 by reduction of its LIM domain at Cys8 (−5%) could lead to upregulation of NF-κB signaling, thereby increasing CD4^+^ T cell activity.

Conversely, H_2_O_2_ might inhibit Vascular endothelial zinc-finger 1 (VEZF1) which can mediate Interleukin-3 (IL-3) transcription (156). IL-3 is known to be produced by CD4^+^ T cells and to activate other T cells and mast cells (157,158). Therefore, loss of zinc-finger binding through reduction of Cys176 and Cys179 (−5%) zinc-finger might downregulate IL-3-dependent T cell activation. Again, suggesting that mtROS plays a role in fine-tuning T cell function.

An advantage of utilizing PMA stimulation was it allowed to uniquely identify mtROS-dependent proteins. However, considering zinc-binding was found enriched as H_2_O_2_ sensitive it would now be interesting to further study the interplay between mtROS second messenger signaling affecting both zinc and calcium signaling. It is known that several protein families such as S100 family (159) can bind both, and it is tempting to speculate that H_2_O_2_ and Ca^2+^ cross-talk will lead to more novel processes that are mtROS sensitive.

Taken together, this work revealed a highly specific redox response in stimulated primary CD4^+^ T cells complementing our understanding on how the mitochondria act as a signaling organelle and how H_2_O_2_ acts as a signaling molecule under redox homeostasis. Inside the mitochondria we have elucidated the potential key regulators in fine-tuning mtROS so that they may mediate their physiologically relevant signals. Additionally, we have provided compelling evidence that the mitochondrial microenvironments likely play a major role for constituting mtROS-dependent protein networks such as the local actin cytoskeleton. Furthermore, we have provided a repository of novel thiol-switch proteins which characterize the mtROS-dependent CD4^+^ T cell immune response. Finally, we propose zinc-finger proteins to constitute a novel H_2_O_2_-mediated transcription/translation regulatory mechanism in which H_2_O_2_ may fine-tune downstream effects on T cell phenotypes in healthy individuals as well as in multiple immune-related diseases.

## Acknowledgments

We acknowledge the work of Reiner Munder in isolating PBMC fractions from whole blood and Dr. Lothar Gröbe for support in T cell isolation by FACS. This work was supported by grants from the Deutsche Forschungsgemeinschaft (PROCOMPAS graduate school, GRK 2223/1 and CRC 854).

## Supplementary Figures

**Supplementary Figure 1:**
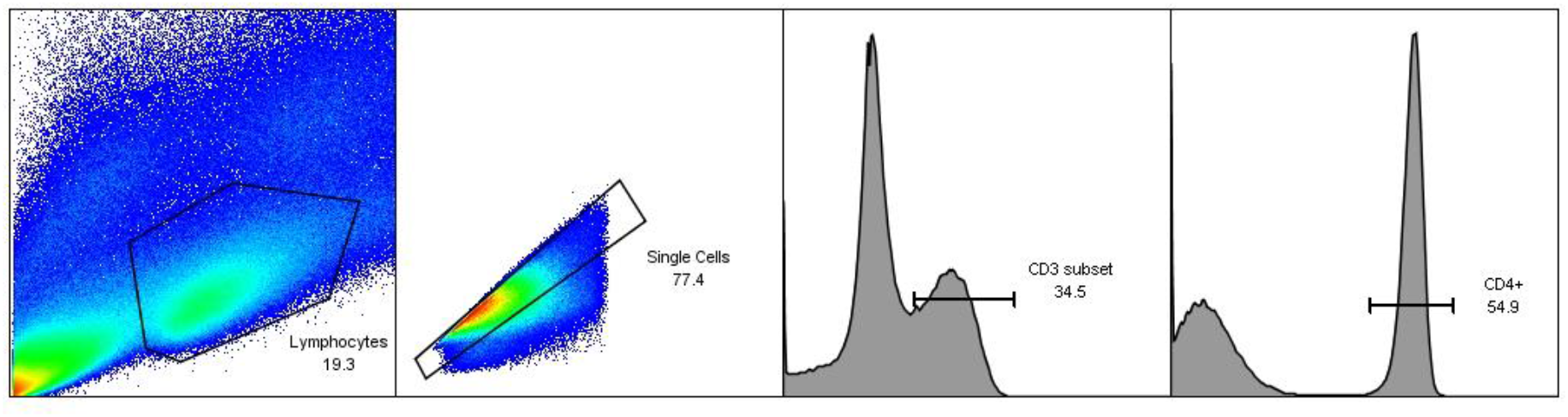
Gating strategy for isolating CD3+CD4+ primary human T cells. PBMCs were isolated from healthy donors and labelled with antibodies directed against CD3, CD4 and CD8. CD3^+^CD4^+^ T cells were sorted by subsequent gating on (1) lymphocytes, (2) single cells, (3) CD3^+^ cells and (4) CD4^+^ cells. CD3^+^CD8^+^ T cells were sorted by subsequent gating on (1) lymphocytes, (2) single cells, (3) CD3^+^ cells and (4) CD8^+^ cells (not depicted).

**Supplementary Figure 2:**
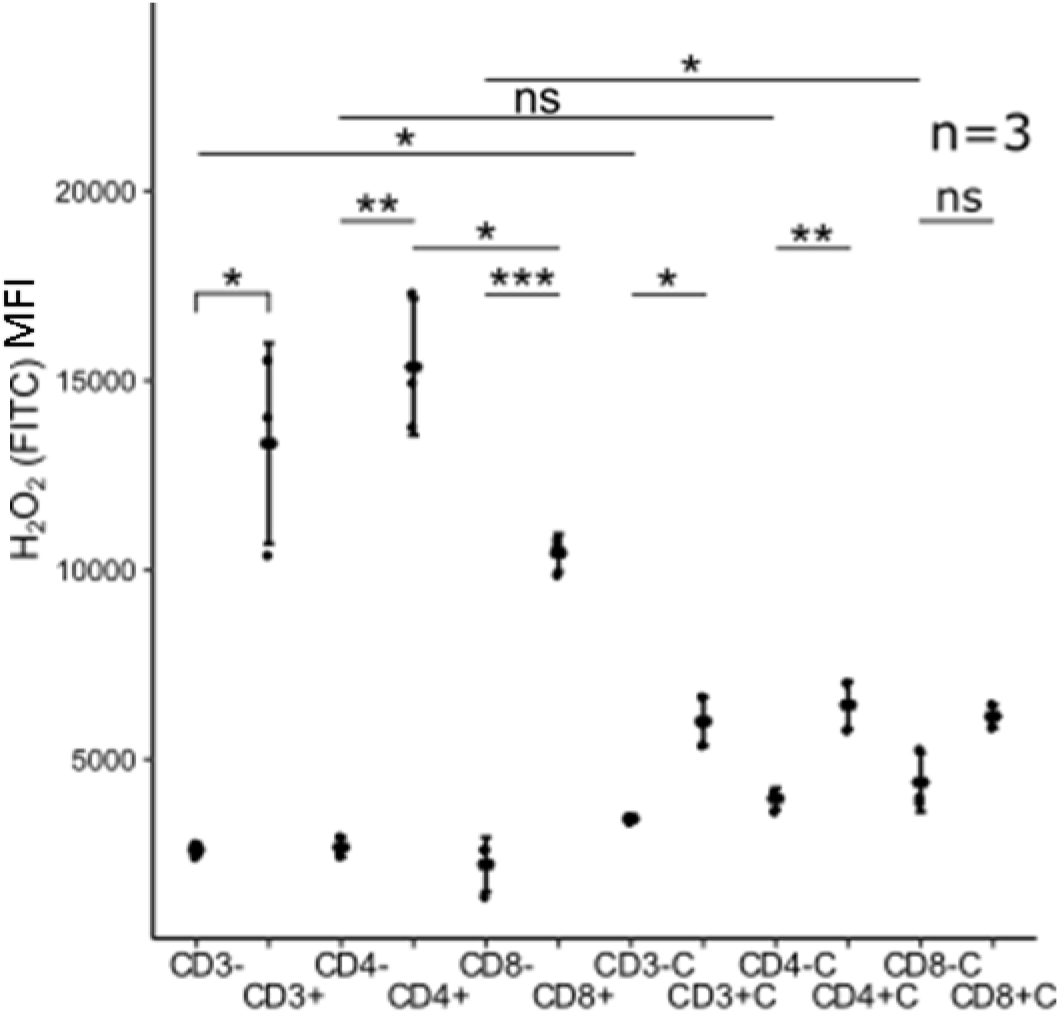
Differential H_2_O_2_ induction in CD3^+^CD4^+^ and CD3^+^CD8^+^ T cells. PBMCs were stimulated with 10 ng/ml PMA for 1 hour (+) in the presence or absence of catalase (C), and intracellular H_2_O_2_ was analyzed in CD3^+^ T cells (CD3), CD3^+^CD4^+^ T cells (CD4) and CD3^+^CD8^+^ T cells (CD8) by measuring H_2_DCFDA using flow cytometry (MFI). Non-stimulated cells (-) were used are controls. Statistical analysis was performed using a two-sided, paired T-test, * p<0.05, ** p<0.01, n=3.

**Supplementary Figure 3:**
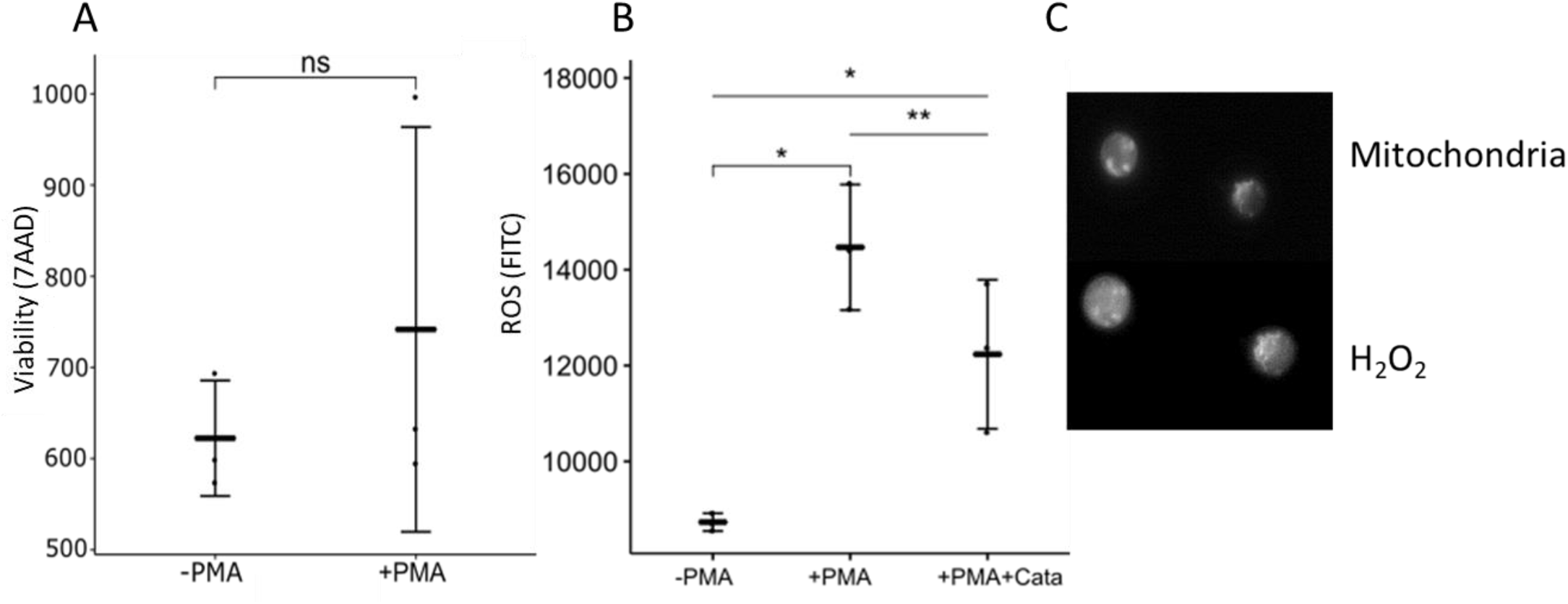
Stress response, H_2_O_2_ induction and mtROS confirmation in CD4^+^ T cells. Isolated CD3^+^CD4^+^ cells were activated with 10 ng/ml PMA for 1 hour (A-C) and analyzed for T cell stress response as seen by 7AAD fluorescence (A), and H_2_O_2_ production as seen by H_2_DCFDA fluorescence in the presence and absence of catalase (B). (C) Fluorescence microscopy images showing co-localization of mitochondria (upper panel) and intracellular H_2_O_2_ (lower panels). Statistical analysis was performed using a two-sided, paired T-test, * p<0.05, ** p<0.01, n=3.

**Supplementary Figure 4:**
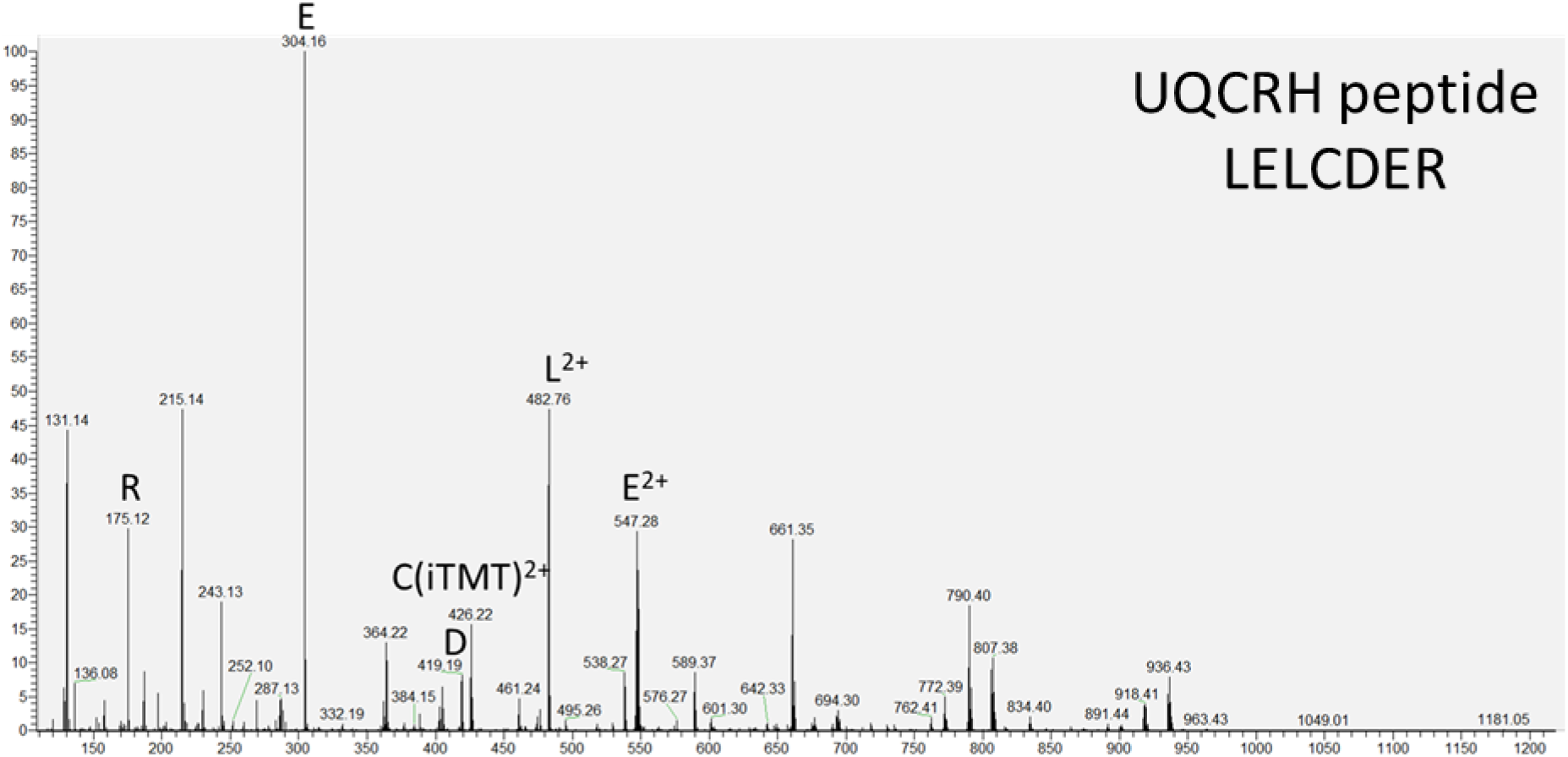
Representative MS^2^ spectra of an iodoTMT-labeled peptide of UQCRH. MS^2^ spectrum showing ion intensities (y-axis) over retention time (x-axis). Intensities peaks for Arginine (R), Glutamate (E), Aspartame (D), IodoTMT-labeled cysteine doubly charged ion (CiTMT2+), Leucine (L) and Glutamate doubly charged ion (E2+) are marked.

**Supplementary Figure 5:**
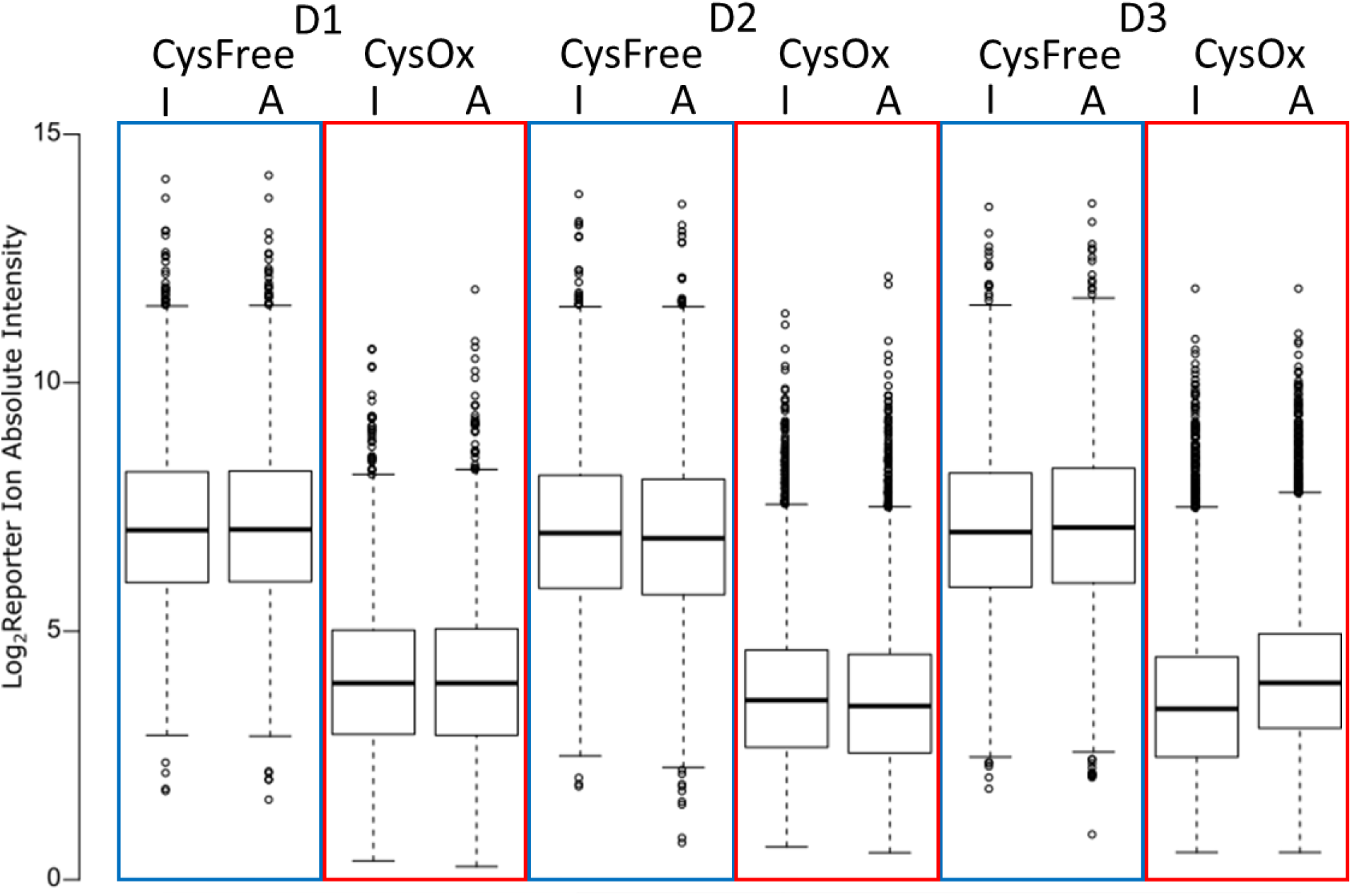
Absolute intensities of each label used to quantify the oxidation state of all cysteine-containing peptides. Distribution of absolute reporter ion intensities (in Log_2_) of each label as used in our multiplex thiol-specific workflow (Figure 2) revealed a general oxidation status of cysteine-containing peptides in non-induced (I) and PMA-induced (A) CD3^+^ T cells from 3 independent donors (D1-D3). Note that the absolute intensities of oxidized peptides (CysOx, red boxes) is lower than the absolute intensities of reduced peptides (CysFree, blue boxes).

**Supplementary Figure 6:**
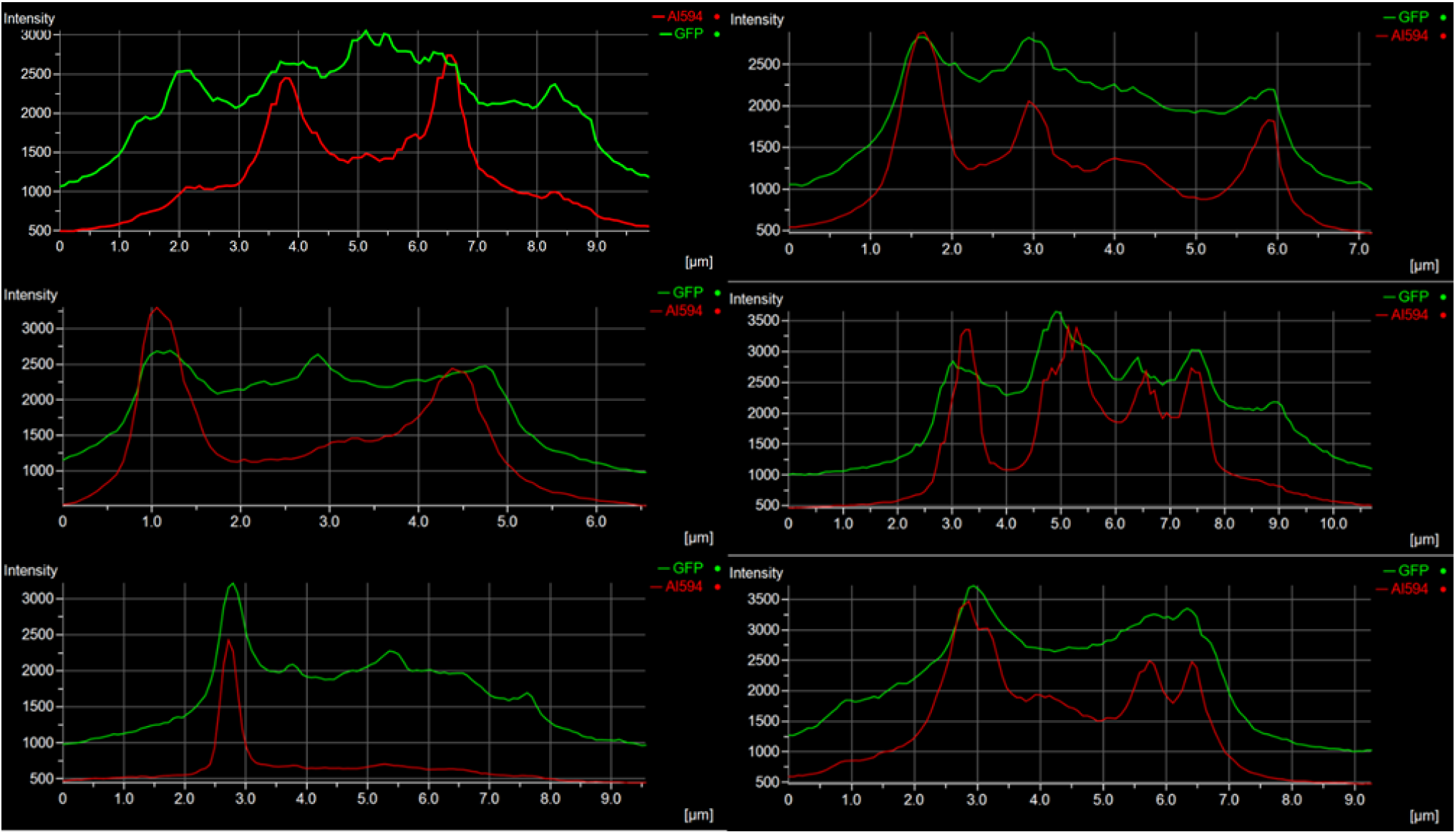
Free zinc (Zn^2+^) and mitochondria intensity profiles showing co-localization in 6 representative human CD3^+^ T cells. PBMCs were stimulated and imaged as described in Figure 6. Representative fluorescence intensity plots of 6 single human CD3^+^ T cells revealing local co-emerging of free zinc (green line) at mitochondria (red line).

